# Multiomic Profiling of Human Clonal Hematopoiesis Reveals Genotype and Cell-Specific Inflammatory Pathway Activation

**DOI:** 10.1101/2022.12.01.518580

**Authors:** J. Brett Heimlich, Pawan Bhat, Alyssa C. Parker, Matthew T. Jenkins, Caitlyn Vlasschaert, Jessica Ulloa, Joseph C. Van Amburg, Chad R. Potts, Sydney Olson, Alexander J. Silver, Ayesha Ahmad, Brian Sharber, Donovan Brown, Ningning Hu, Peter van Galen, Michael R. Savona, Alexander G. Bick, P. Brent Ferrell

## Abstract

Clonal hematopoiesis (CH) is an age-associated phenomenon that increases risk for hematologic malignancy and cardiovascular disease. CH is thought to enhance disease risk through inflammation in the peripheral blood^1^. Here, we profile peripheral blood gene expression in 66,968 single cells from a cohort of 17 CH patients and 7 controls. Using a novel mitochondrial DNA barcoding approach, we were able to identify and separately compare mutant *TET2* and *DNMT3A* cells to non-mutant counterparts. We discovered the vast majority of mutated cells were in the myeloid compartment. Additionally, patients harboring *DNMT3A* and *TET2* CH mutations possessed a pro-inflammatory profile in CD14+ monocytes through previously unrecognized pathways such as galectin and macrophage Inhibitory Factor (MIF). We also found that T cells from CH patients, though mostly un-mutated, had decreased expression of GTPase of the immunity associated protein (GIMAP) genes, which are critical to T cell development, suggesting that CH may impair T cell function.

**Key points:**

- CD14+ monocytes from clonal hematopoiesis patients stimulate inflammation through increased cytokine expression.
- T cells from clonal hematopoiesis are deficient in GIMAP expression, suggesting CH may impair T cell differentiation.

## Introduction

With age, hematopoietic stem cells acquire mutations in driver genes such as DNA Methyltransferase 3A (*DNMT3A)* and Tet Methylcytosine Dioxygenase 2 (*TET2),* resulting in a selective advantage and clonal hematopoiesis. CH is a risk factor not only for hematologic malignancy but also for multiple diseases of aging including cardiovascular disease, kidney disease, and osteoporosis^1–5^. While many epidemiological analyses consider CH as a single entity, the literature reveals that significant associations are gene specific. For example, *TET2* is more strongly associated with a pro-inflammatory disease mechanism across multiple forms of cardiovascular disease^1,6^ while *DNMT3A* CH is associated with heart failure^7,8^ and osteoporosis^5^.

Much attention has focused on how CH mutations lead to skewed hematopoiesis in hematopoietic stem and progenitor cells (HSPC)^9,10^. However, little attention has focused on the peripheral compartment. Circulating immune cells with CH mutations are morphologically and immunophenotypically similar to their non-mutated counterparts, making direct comparisons difficult in primary human tissues. Whether primary cell intrinsic transcriptional changes or secondary microenvironment effects, or both, drive pathological phenotypes is unknown.

Though transcriptional profiling of single cells has become routine, it remains difficult to extract genotype and transcriptional data out of the same cell. Since *DNMT3A* and *TET2* CH blood samples are a mixture of mutated and non-mutated cells, both genotyping and transcriptomic sequencing modalities are necessary to delineate these cells. Cell-intrinsic consequences may arise directly from the somatic mutation while extrinsic, indirect consequences may arise from altered cell-cell interactions or secreted immune effectors. These phenomena could be distinguished by identifying mutant and non-mutant cells from the same sample. Several technologies have sought to close this gap by selectively amplifying the mRNA transcriptome and using this to genotype cells^11–14^. This approach is effective in HSPCs that express *DNMT3A* and *TET2*; however, genotyping is less efficient in cells that do not express these genes, such as fully differentiated cells in the peripheral blood^12,13,15^.

To overcome this, we combined single-cell RNA-sequencing (scRNA-seq) with cell-specific mitochondrial DNA barcoding to simultaneously resolve single-cell DNA mutation status for 100% of cells^15^. Our analysis of 66,968 single cells from 17 individuals with *TET2* CH or *DNMT3A* CH and 7 age-matched controls finds novel mechanisms of CH-driven inflammation and enables direct comparison between peripheral CH mutated and wildtype cells across individuals.

## Methods

### Primary patient samples

All patients in this study consented to all study procedures under VUMC institutional review board approved research protocols (IRBs #210022, #201583) in accordance with the Treaty of Helsinki. Adult patients able to give consent were recruited from VUMC clinics who had known CH mutations as a result of clinical evaluation or patients who were at risk of having a CH mutation. All patients were confirmed to be without active hematological malignancy at the time of enrollment. Fresh PBMCs were isolated using Ficoll separation. Following low-speed centrifugation, pelleted cells were resuspended in freezing media (88% FBS and 12% dimethyl sulfoxide (DMSO)) and placed in liquid nitrogen.

### DNA extraction and CH variant calling

All enrolled patients underwent targeted sequencing to evaluate for the presence of CH mutations. DNA was extracted using Qiagen Mini kits Cat #27104 according to manufacturer’s recommendations. We sequenced samples using a custom capture panel designed to tile known CH genes, targeting 600x read depth coverage as previously described^16^. Somatic mutations were called using publicly available methods in workflow description language in Mutect2 on the Terra Platform (https://terra.bio/). A putative variant list was formulated and then cross referenced with a list of known CH driver mutations^1^. Variants were then filtered for read quality including sequencing depth and minimum alternate allele read depth.

### scRNAseq library preparation

Cryopreserved PBMCs from CH patients and controls were thawed at 37 °C and washed with complete RPMI (RPMI + 10%FBS + 1%PS, cRPMI) to remove freezing media. PBMCs from each sample (500,000 cells) were plated. Cells were pooled following staining with unique hashtag antibody oligonucleotide-conjugates (HTO) (30 minutes) staining (Biolegend, TotalSeq-B). Pooled samples were immediately run on a 10x Chromium Controller after preparation with a 10X Chromium 3’ library preparation kit (10X Genomics) to create scRNA-seq libraries.

### 10X single-cell sequencing and data preparation

Next-generation 150-nt paired-end sequencing was conducted on an Illumina Novaseq6000 using the cDNA libraries produced by the 10X Chromium library (**Supplemental Table6**). CellRanger Count (10X Genomics) was used to filter low quality reads and align to the GRCh38 reference genome using STAR as described elsewhere^17^. Resulting matrices from the CellRanger pipeline were then converted into Seurat^18,19^ objects. Demultiplexing was performed using the HTODemux function in the R package Seurat, applying a positive quantile value of 0.99. Cells containing 15% or more reads mapping to the mitochondrial genome were filtered. Similarly, we filtered cells with less than 250 genes and 500 UMIs respectively. Remaining doublets were removed with the R package DoubletFinder^20^, using the first 10 principal components and a doublet formation rate of 7.5%^21^. One lane had an abnormally high number of doublets, so a more stringent filter was applied for that lane using the DoubletFinder metric pANN of 0.28. Batch correction was performed with the R package Harmony (v0.1.1). R package Harmony (v0.1.1).

After performing dimensionality reduction with the function RunUMAP from Seurat and calculating clusters with the function FindClusters (resolution = 0.75), cell type assignment was performed using ScType (v1.0). Low confidence cell types were annotated manually. Mitochondrial and ribosomal genes were removed.

### Single-cell mitochondrial enrichment

Mitochondrial enrichment of 10x Genomics v3 3’ cDNA was performed using primer sequences as described in Miller, TE et al^22^. Briefly, 10X 3’ cDNA was amplified using primers encompassing the entire mitochondrial genome along with i5 indexes (**Supplemental Table 5)**. The samples were pooled, mixed, and incubated with 1.0X AMPure XP Beads. Following incubation, i7 indexes were added. The amplified cDNA library was purified with AMPure XP beads and eluted in TE Buffer.

### Single-cell mitochondrial DNA sequencing, read processing, and variant calling

MT-DNA processing and variant calling were carried out as previously published^15^. Briefly, fastq files from MT-DNA enrichment were filtered for reads associated with low-frequency cell barcodes (CB) and trimmed to remove the UMI and CB. Reads were aligned using STAR to hg38. Next, we used maegatk to call variants across the mitochondrial genome^15,23^. Maegatk calls mtDNA variants using combined CB from both scRNAseq and MAESTER enrichment for variants with at least five supporting reads.

### Identification and selection of informative single cell mtDNA variants

Using output from Maegatk, an allele frequency matrix was constructed using all possible variants in the mitochondrial genome. Cell type annotations were transferred using the cell barcodes from the RNA expression dataset. A table of MT variants was constructed based on cell type enrichment for individual MT mutations. MT mutations enriched within monocytes and absent or very low in lymphoid lineage cells were considered candidate markers of CH mutations. Single-cell DNA sequencing was performed with Tapestri scDNAseq and colony based single cell DNA sequencing to confirm candidate MT mutations alignment with CH mutations.

### Single-cell DNA sequencing via MissionBio Tapestri

Single-cell DNA sequencing was performed using the Tapestri platform. Cells were stained with the BioLegend Total-Seq D Heme Oncology Panel and the Human TruStain FcX antibody. Targeted DNA amplification was carried out using custom designed probes from MissionBio (**Supplemental Table 8**). The amplified DNA was released from individual oil droplets using Ampure XP beads. The final product was quantified using a Qubit fluorometer from ThermoFisher and assessed for quality on an Agilent Bioanalyzer. Samples were pooled prior to sequencing with a 25% spike-in of PhiX and run on a NovaSeq 6000 S4 flow cell from Illumina to generate 150 bp paired-end reads. Sequencing was performed at the Vanderbilt Technologies for Advanced Genomics (VANTAGE) sequencing core.

### Pipeline processing and variant filtering for Tapestri single-cell DNA sequencing

Single-cell DNA samples were processed using the Tapestri Pipeline v1.8.4. Adapters were trimmed and reads were aligned to the hg19 reference genome. Variants were called using GATK 3.7 and filtered based on quality scores, read depth, and genotype frequency. Informative variants were annotated, and cells were clustered based on their genotypes.

To annotate cell populations, unsupervised hierarchical clustering was performed on the antibody-oligo conjugate (AOC) data. Reads were normalized and AOCs with low expression were removed. Principal component analysis was conducted on the normalized AOCs, and the first ten principal components were used for UMAP coordinate calculation. The resulting cells were clustered using a k state of 100, and clusters with noisy AOC expression were eliminated. The remaining clusters were annotated based on expert knowledge of surface marker expression.

### Differential expression analysis and pathway analysis

Differential gene expression was calculated using a pseudobulk-like approach in which measurements from groups of similar cells were summed. Cells were separated by genotype and cell type then clustered using the Python module Metacell-2 (v0.8.0). We followed standard procedures from the Metacell-2 vignette. We excluded gene modules that had an average correlation with cell cycling genes of 0.75 or greater. Differential expression was calculated for genes that had at least 10 transcripts in at least 85% of metacells using the R package DESeq2. We performed Wald tests of significance with Benjamini-Hochberg multiple testing correction. Genes that were sex-specific and red-blood-cell-specific were removed. Pathway analysis was performed on differential expression results using the function gseGO with ont = “ALL”, minGSSize = 50, maxGSSize = 800, nPermSimple = 10000 from the R package clusterProfiler (v4.8.1).

Cell signaling interactions were predicted from single-cell RNA sequencing data with the R package CellChat (v1.6.1). Genes with extremely high or low expression were removed with the parameter trim = 0.1. Comparisons were restricted to cell types that had at least 25 cells with the parameter min.cells = 25.

To compare mutant and wildtype monocyte cell states, we performed pairwise DGE analysis by using the MAST method with false-discovery rate (FDR) correction. To reduce transcriptional noise prior to DGE, we only included genes that were detected in at least 10 cells. We then applied the Hurdle model from the MAST R package (v.1.24.1) and adjusted for the cellular detection rate to determine significant differences in gene expression (threshold: absolute value of the log fold-change coefficients > 0.25, FDR > 0.05).

### Phospho-specific flow cytometry

Cryovials of cryopreserved cells from healthy donors and CH patients were thawed and washed with 10 mL of cRPMI. The cells were stained for viability with AlexaFlour700 (Invitrogen, cat#P10163) and counted. An aliquot of 500,000 cells were plated in 200 µL of media in a 96-well plate and stimulated with 20 ng/mL of IL-6 (Peprotech) for 15 minutes. Cells were then fixed with 1.6% PFA at room temperature and permeabilized with 150 µL of methanol at -80C for at least 30 minutes. Cells were resuspended in 180 µL of PBS and fluorescence cell barcoding performed as previously described^24^ with serial dilutions of Pacific Blue (LifeTechnologies, cat#P10163, PB) and Pacific Orange (Invitrogen, cat#P30253, PO) dyes for 30 minutes in the dark at RT. Two concentrations of PB were prepared (20 and 4 µg/mL), while six concentrations of PO were prepared (7.00, 2.99, 1.27, 0.54, 0.23, 0.10 µg/mL). Barcoding was quenched with 80 µL of cell staining media. The barcoded cells were then collected into a single tube and stained with a cocktail of antibodies: CD33 PECy7 (5 µL per 100 µL stain, Biolegend, cat#303434, clone WM53) and pSTAT3 AlexFlour488 (2.5 µL per 100 µL stain, Biolegend, cat#651006, clone 13A3-1) for 30 minutes prior to acquisition on a BD 5-laser Fortessa flow cytometer.

## Results

We used scRNA-seq with cell-specific mitochondrial DNA sequencing to resolve single-cell genomic DNA mutation status and investigate pathological mechanisms of CH (**Fig. 1A, methods**). Peripheral blood mononuclear cells (PBMCs) from 8 *TET2*, 9 *DNMT3A* and 7 age-matched controls (ages 47-89) were selected from a prospective CH observational study which was designed to capture patients at high risk for CH through a robust referral network (**Fig. 1A-B** **and Table 1**).

**Fig. 1:**
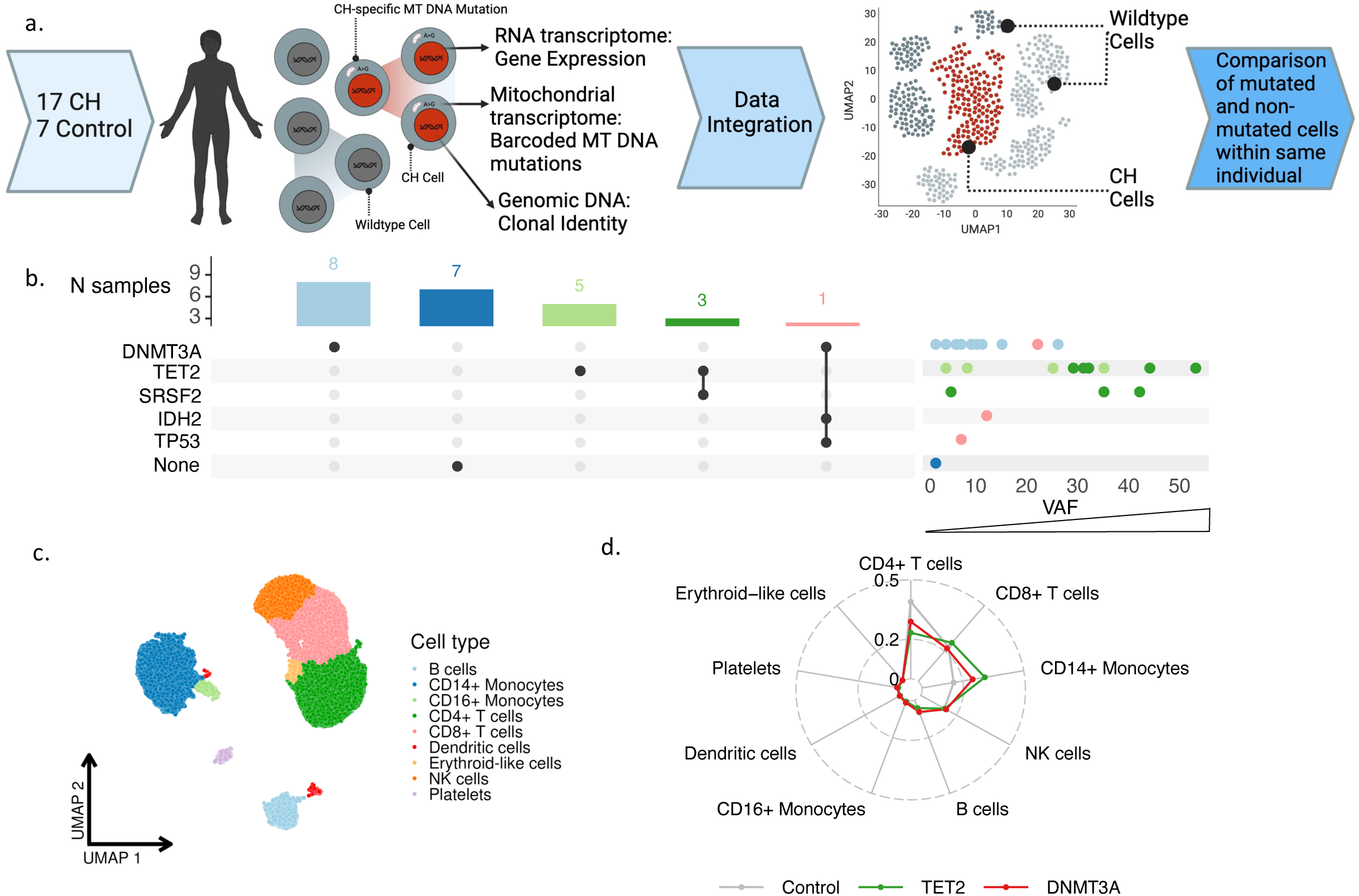
Single-cell RNA sequencing reveals distinct cell type profiles in clonal hematopoiesis. **A)** Cells from 17 CH patients and 7 controls were processed via scRNAseq. Mutational lineage tracing was performed to link genomic variants to scRNAseq results. **B)** UpSet plot displaying CH mutations for each patient with accompanying dot plot showing variant allele frequency for each mutation. Color corresponds to mutational group. Dots correspond to mutations, so patients with multiple CH mutations have separate dots for each mutation. **C)** UMAP displaying cell type clusters, as defined by unsupervised clustering. Clusters were annotated using scType. **D)** Radar plot showing cell type proportions for controls, patients with *TET2* mutations, and patients with *DNMT3A* mutations.

**Table 1.** Demographic features of CH and control patients. Values are listed by counts and by percentages for categorical variables and by mean and standard deviation for continuous variables.

|  | **Control**<br>**(N=7)** | **DNMT3A**<br>**(N=9)** | **TET2**<br>**(N=8)** |
| --- | --- | --- | --- |
| **Sex** |  |  |  |
| Female | 4 (57.1%) | 2 (22.2%) | 2 (25.0%) |
| Male | 3 (42.9%) | 7 (77.8%) | 6 (75.0%) |
| **Race** |  |  |  |
| White | 7 (100%) | 9 (100%) | 6 (75.0%) |
| Asian | 0 (0%) | 0 (0%) | 1 (12.5%) |
| Missing | 0 (0%) | 0 (0%) | 1 (12.5%) |
| **Age** |  |  |  |
| Mean (SD) | 75 (14) | 67 (15) | 75 (8.8) |
| Missing | 1 (14.3%) | 0 (0%) | 0 (0%) |
| **Hx tobacco use** | 3 (42.9%) | 5 (55.6%) | 3 (37.5%) |
| **Hx CAD** | 3 (42.9%) | 3 (33.3%) | 2 (25.0%) |
| **Hx CHF** | 1 (14.3%) | 1 (11.1%) | 1 (12.5%) |
| **Systolic BP** |  |  |  |
| Mean (SD) | 130 (14) | 120 (16) | 130 (11) |
| **Diastolic BP** |  |  |  |
| Mean (SD) | 73 (16) | 70 (9.1) | 74 (11) |
| **Heart rate** |  |  |  |
| Mean (SD) | 70 (11) | 78 (14) | 68 (9.4) |
| Missing | 0 (0%) | 1 (11.1%) | 0 (0%) |

To trace the effects of CH mutations on peripheral blood cell type proportions, we derived cell type annotations based on known marker genes (**Fig. 1C-D, Supplemental Fig. 1).** There were no significant differences in cell type proportions on routine clinical laboratories **(Supplemental Table 1**). Notably, four patients had multiple CH mutations with concomitant cytopenias without bone marrow dysplasia, meeting diagnostic criteria for clonal cytopenias of undetermined significance (CCUS)^25^.

To annotate mutant and non-mutant cells from the same sample, we combined single-cell targeted amplicon sequencing (scDNAseq) and 3’ RNA mitochondrial lineage tracing in our *TET2* and *DNMT3A* patients (**Fig. 2A**). PBMCs from both *TET2* and *DNMT3A* patients were processed through the scDNAseq pipeline (Mission Bio) which captures known genomic CHIP mutations and co-occurring mitochondrial variants. We also profiled the immunophenotype of the samples by combining scDNAseq with oligo-conjugated antibodies to annotate cell populations **(Fig 2B, 2E, and Supplemental Fig. 2-B).** In one patient with a known *TET2* mutation at chr4:106157967 with 51% VAF **(Supplemental Fig 2, Supplemental Table 2)** our scDNAseq analysis revealed a single mitochondrial variant (MT 7754G>C) that was concordant with cells harboring the known *TET2* mutation, suggesting common lineage. We found 492 cells that carried both the *TET2* mutation and the MT 7754G>C variant and 492 cells that carried neither variant. We excluded a marginal number (n = 15) of cells where only the 7754G>C variant was detected. The mature myeloid cell compartment was heavily enriched for both the *TET2* mutation and the mitochondrial variant **(Fig. 2C-D, Extended Data** Fig. 3A). We repeated the analysis for a sample with a known *DNMT3A* mutation with 24% VAF **(Fig 2F-G, Supplemental Fig 2)**. The concomitant genomic and mitochondrial variants (chr2:25470560 and MT 747A>G) were detected in most myeloid cells and a moderate proportion of lymphocytes, consistent with previous knowledge regarding *DNMT3A* mutations in hematopoiesis^26^ (**Fig. 2F-G** **and Extended Data** Fig. 3B). There were 111 *DNMT3A* cells where the 747A>G variant was not detected, indicating that 747A>G marks a cell population that is subclonal to *DNMT3A*. We validated the co-occurrence of the CH variant and 747A>G using primary template amplification of genomic DNA from single cell colonies (**Extended Data** Fig. 3D**, see Methods**). Subsequently, we were able to use the MT DNA single nucleotide variant (SNV) as an identifying ‘barcode’ for the mutant cell annotation, allowing for the partition of mutant and wildtype populations within our scRNAseq data (**Fig. 2H-K**). In the corresponding single-cell RNA dataset evaluating for the *TET2* sample, we found a significant myeloid bias among cells identified with MT 7754G>C (log2(fold change) = 4.146, FDR = 0.001, **Fig 2H-I**). In DNMT3A scRNAseq sample, we identified cells with 747A>G, finding a less severe monocytic skew accounting for the lower relative VAF compared to the *TET2* sample (**Fig. 2J-K**).

**Fig. 2:**
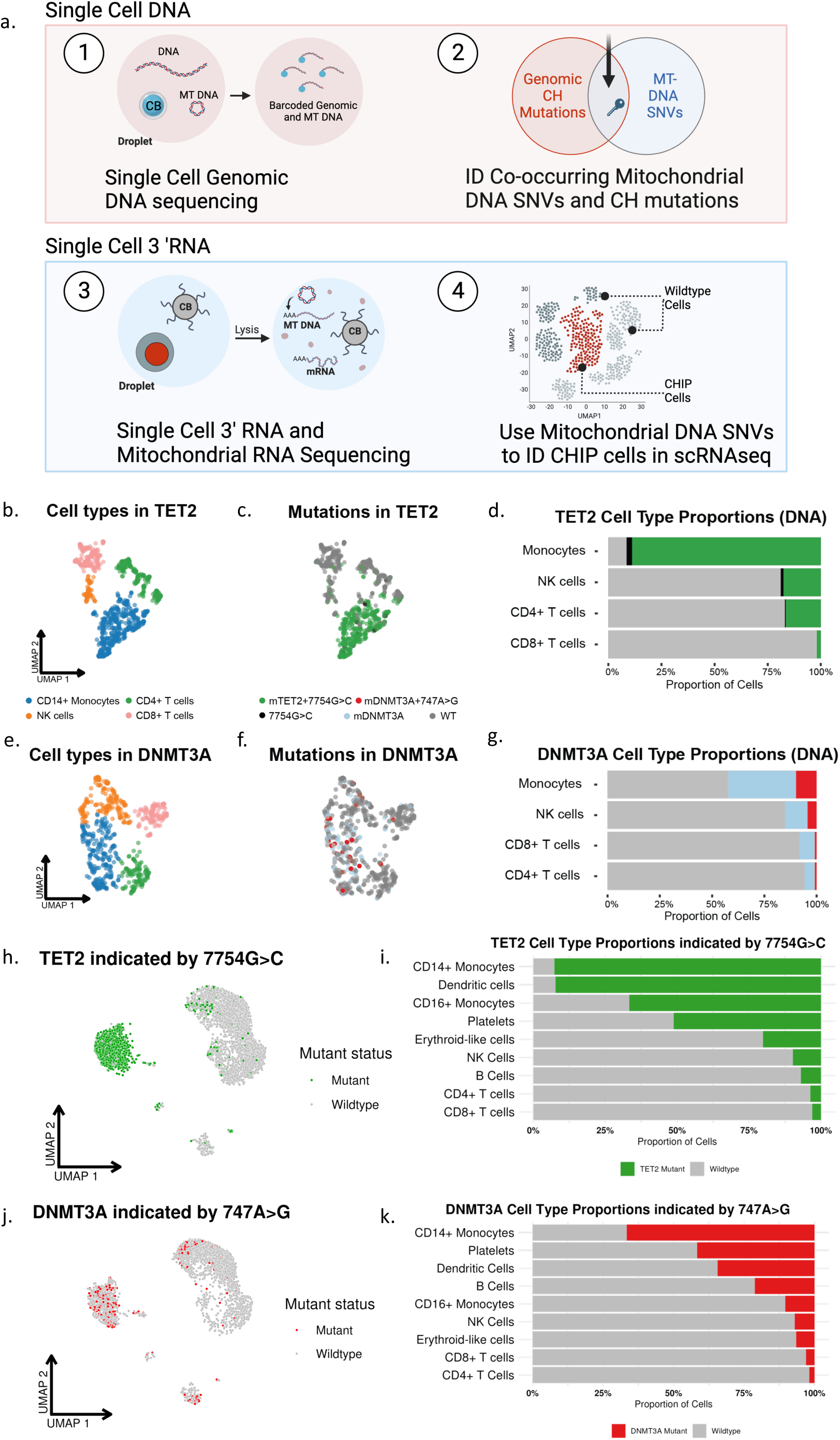
Single-cell DNA and RNA sequencing define cells carrying CH mutations. **A)** Schematic showing experimental design. **B)** UMAP showing cell types for cells from *TET2* patient CH-21-014. **C)** UMAP showing mutational status for cells from *TET2* patient CH-21-014. **D)** Stacked bar plot quantifying proportion of mutant cells per cell type for cells from *TET2* patient CH-21-014. **E)** UMAP showing cell types for cells from *DNMT3A* patient CH-20-046. **F)** UMAP showing mutational status for cells from *DNMT3A* patient CH-20-046. **G)** Stacked bar plot quantifying proportion of mutant cells per cell type for cells from *DNMT3A* patient CH-20-046. **H)** UMAP showing predicted mutational status for cells from *TET2* patient CH-21-014, based on presence of MT mutation 7754G>C. **I)** Stacked bar plot quantifying proportion of mutant cells per cell type for cells from *TET2* patient CH-21-014. **J)** UMAP showing predicted mutational status for cells from *DNMT3A* patient CH-20-046, based on presence of MT mutation 747A>G. **K)** Stacked bar plot quantifying proportion of mutant cells per cell types for cells from *DNMT3A* patient CH-20-046.

We applied our mitochondrial lineage tracing *method* to a total of 4 *TET2* and 2 *DNMT3A* patients to identify CH clones (**Supplemental Fig. 3).** We first performed differential gene expression testing (DGE) on CD14+ monocytes comparing CH mutant cells to their wildtype (WT) counterparts. This resulted in 70 differentially expressed genes in *TET2* mutants whereas there were zero differentially expressed genes in *DNMT3A* mutants compared to WT counterparts. We then evaluated mutant *TET2* and *DNMT3A* CD14+ monocytes against unaffected control CD14+ monocytes. We identified 202 and 122 differentially expressed genes (DEGs) in *TET2* and *DNMT3A* CD14+ monocytes, respectively (**Supplemental Table 7)**. There were 12 overlapping DEGs when comparing mutant *TET2* CD14+ monocytes to WT cells and when comparing to controls (**Supplemental Fig. 3**). Notable among these were inflammatory mediators *CXCL1, CXCL3, and IL1B* (False Discovery Rate [FDR] < 1 x 10^-20^ for all comparisons) **(Fig. 3A, Supplemental Table 7**). Top DE genes in mutant *DNMT3A* monocytes compared to controls included C-C Motif Chemokine Ligand 4 (*CCL4*, FDR = 1.26 x 10^-4^), C-C Motif Chemokine Ligand 2 (*CCL2,* FDR = 2.78 x 10^-12^), and C-C Motif Chemokine Ligand 7 (CCL7, FDR = 1.34 x 10^-7^) (**Fig. 3B, Supplemental Table 7**). Pathway analysis showed upregulation of leukocyte activation and cell adhesion in mutant *TET2* monocytes whereas mutant *DNMT3A* monocytes had enrichment in regulation of cellular death pathways and leukocyte migration (**Fig. 3C-D**).

**Fig. 3:**
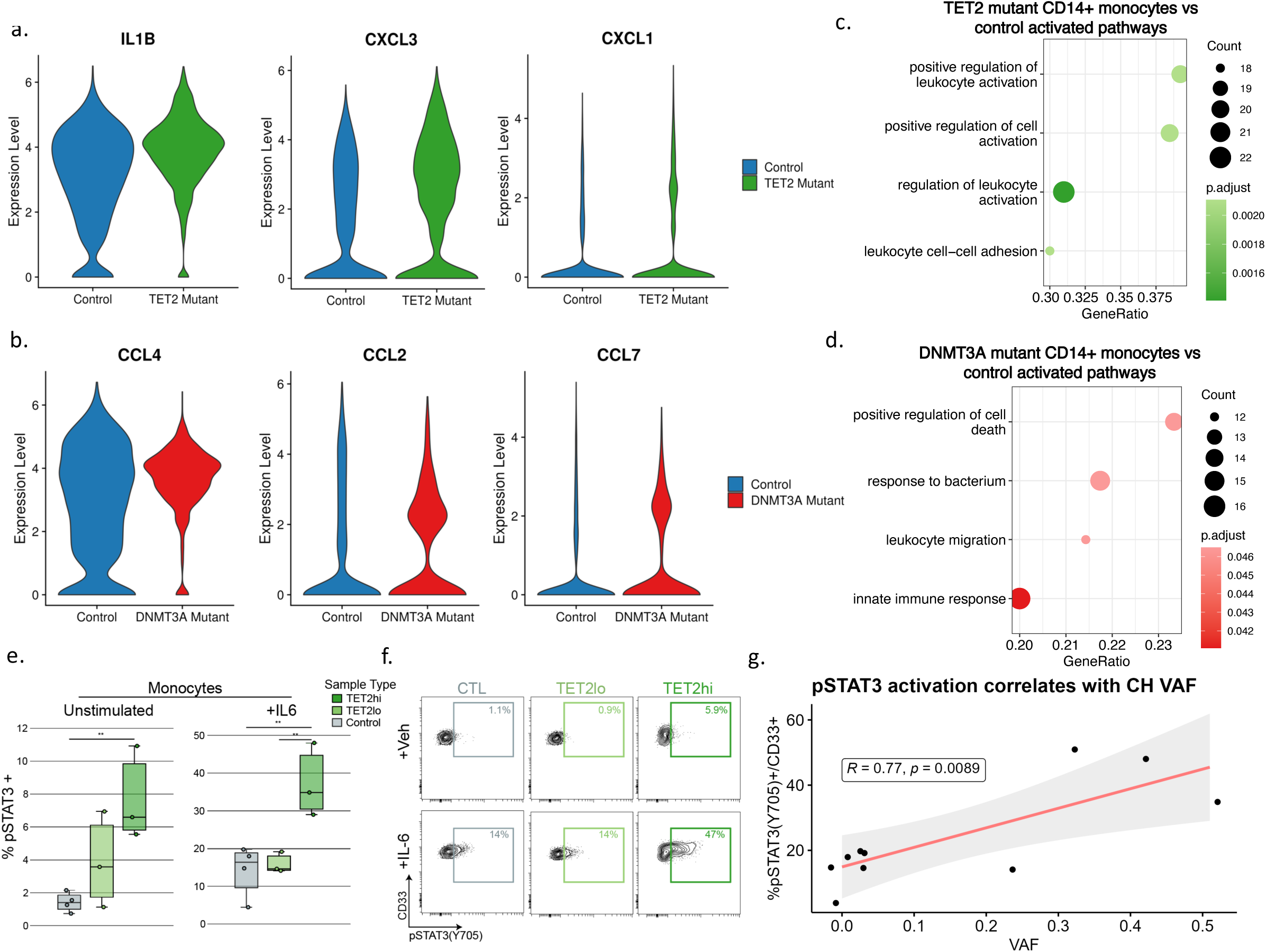
CD14+ monocyte mutant vs control analysis highlights inflammatory cell-intrinsic effects of CH mutations. **A)** Violin plots displaying expression of cytokines in control CD14+ monocytes and *TET2* mutant CD14+ monocytes. **B)** Violin plots displaying expression of cytokines in control CD14+ monocytes and *DNMT3A* mutant monocytes. **C)** GSEA based on results from differential expression analysis comparing *TET2* mutant CD14+ monocytes to control CD14+ monocytes. **D)** GSEA based on results from differential expression analysis comparing *DNMT3A* mutant CD14+ monocytes to control CD14+ monocytes. **E)** Quantification of phospho-flow cytometry displaying pSTAT3(Y705)+ in the CD33+ gate of large VAF *TET2* CH samples (>25%, n = 3) compared to both low VAF (<25%, n = 3) and control samples (n = 4) (** = p < 0.01 by ANOVA with Tukey’s HSD). **F)** Same as in (E), with IL-6 stimulation condition. **G)** Pearson correlation of %pSTAT3(Y705)+ in the CD33+ gate and VAF following IL-6 stimulation. For samples with multiple mutations, the highest VAF value was selected for the analysis.

Noting the increased expression of IL-1B, a prominent downstream mediator of the IL-6 pathway among mutant *TET2* monocytes, we sought to further evaluate whether signaling along this axis was a cell intrinsic or cell extrinsic phenomenon. To do this, we employed phosphospecific flow cytometry to measure response to IL-6 in high VAF *TET2* mutant (*TET2*hi), low VAF *TET2* mutant (*TET2*lo), and controls. The basal pSTAT3+ monocyte percentage was significantly higher in *TET2*hi monocytes compared to controls. All samples showed some response to IL-6, while *TET2*hi monocytes had the highest proportion of pSTAT3+ cells, significantly higher than both control and *TET2*lo samples, in response to IL-6 stimulation (**Fig. 3E-F**). Notably, there was a linear increase in the proportion of pSTAT3+ cells after IL-6 stimulation in accordance with increasing VAF (R = 0.77, p = 0.008), suggesting cell intrinsic altered signaling among the mutant fraction (**Fig. 3G**).

We then queried whether there were also cell-extrinsic effects of CH mutations in our cohort as has been recently reported.^27^ To determine this, we compared grouped RNA expression profiles from CD14+ monocytes between *DNMT3A* or *TET2* and control patients since this would include both mutated and non-mutated cells. To reduce potential for false discovery from high dropout rates, we partitioned our dataset into metacells^28^ prior to performing DGE analysis. The top DEGs among *TET2* patients included fibronectin 1 (*FN1*, adj p = 2.39 x 10^-26^) and Fc Epsilon Receptor II (*FCER2*, adj p = 2.4 x 10^-14^), which encodes CD23. Both of these are important components of monocyte adhesion^29,30^ (**Fig 4A**). This contrasts with the top DEGs from *DNMT3A* CD14+ monocytes which included interferon induced transmembrane protein 2 (*IFITM2*, adj p = 2.37 x 10^-18^) and adhesion G protein-coupled receptor E5 (*ADGRE5*, adj p = 4.4 x 10^-14^), genes involved in monocyte adhesion^31^ and differentiation^32^ (**Fig 4B**). While the specific genes impacted were different between *TET2* and *DNMT3A* comparisons, the pathways they converged on were similar. In general, genes related to immune responses and leukocyte activation were upregulated whereas genes related to transport activity and endoplasmic reticulum regulation were downregulated (**Fig. C-D).** Similarly, GSEA highlighted convergent pathways between *TET2* and *DNMT3A* CD14+ monocytes including leukocyte activation, regulation of leukocyte activation, and regulation of cell activation (**Fig. 2E-F**). Using CellChat^33^ to infer intercellular interactions in our scRNA-seq data, we found CD14+ monocytes from TET samples exhibited enhanced signaling across IL-1, macrophage migration inhibitory factor (*MIF*), and Galectin, all parts of the inflammatory signaling axis (**Fig. 2G**). CD14+ monocytes from *DNMT3A* samples also exhibited enhanced IL-1 and galectin signaling and uniquely had elevated integrin beta 2 (ITGB2) signaling **(Fig 2H, Supplemental Table 7**).

**Fig. 4:**
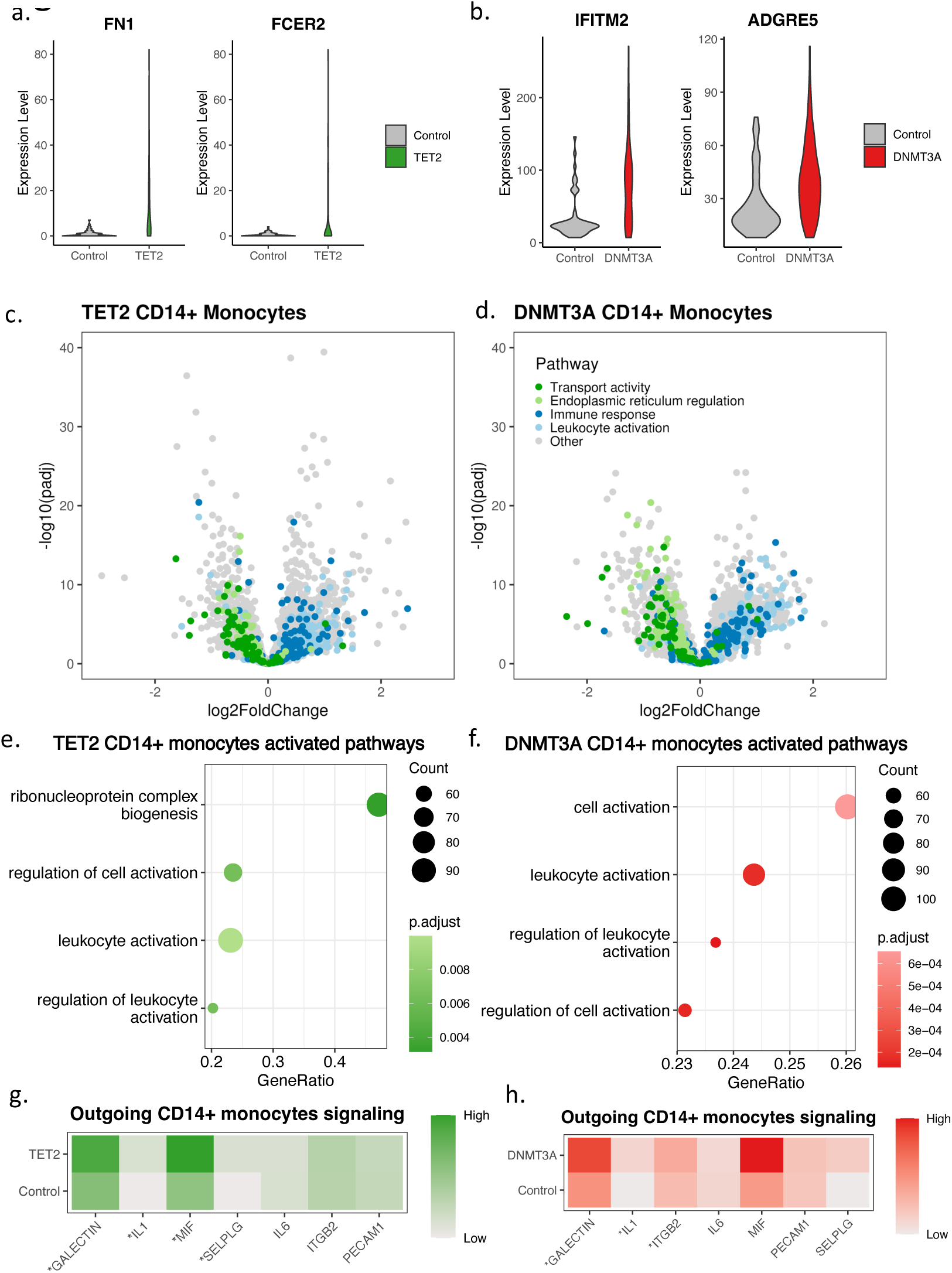
Genotype-grouped CD14+ monocyte vs control comparisons highlight inflammatory cell-extrinsic effects of CH mutations. **A)** Violin plots displaying expression of *FN1* and *FCER2* in CD14+ monocytes from controls and CD14+ monocytes from patients with *TET2* mutations. **B)** Violin plots displaying expression of *IFITM2* and *ADGRE5* in CD14+ monocytes from controls and CD14+ monocytes from patients with *DNMT3A* mutations. **C)** Volcano plot showing results of differential expression analysis comparing CD14+ monocytes from patients with *TET2* mutations to CD14+ monocytes from controls, colored by biological pathway. **D)** Volcano plot showing results of differential expression analysis comparing CD14+ monocytes from patients with *DNMT3A* mutations to CD14+ monocytes from controls, colored by biological pathway. **E)** GSEA based on results from differential expression analysis comparing CD14+ monocytes from patients with *TET2* mutations to CD14+ monocytes from controls. **F)** GSEA based on results from differential expression analysis comparing CD14+ monocytes from patients with *DNMT3A* mutations to CD14+ monocytes from controls. **G)** Heatmap showing predicted outgoing signaling from CD14+ monocytes from patients with *TET2* mutations and from controls for pathways involved in inflammatory response and immune cell migration, as determined by CellChat (* indicates p-value < 0.05/7). **H)** Heatmap showing predicted outgoing signaling from CD14+ monocytes from patients with *DNMT3A* mutations and from controls for pathways involved in inflammatory response and immune cell migration, as determined by CellChat (* indicates p-value < 0.05/7).

When evaluating signaling patterns between cell types, we noted increased signaling from both CH CD14+ monocytes to T cells leading us to investigate the impact of CH on T cells (**Fig. 5A-B**). We found that genes involved in T cell activation and immune response were highly expressed in both *TET2* and *DNMT3A* samples compared to controls (**Fig. 5C-D, Supplemental Table 7)**. Each member of the GTPase of the immune associated nucleotide binding protein (GIMAP) family, which plays a critical role in proper T and B cell differentiation^34,35^ was downregulated in CD4+ T cells and CD8+ T cells in both *TET2* and *DNMT3A* samples (**Fig. 5E-F, Supplemental Fig 4**). *GIMAP1* and *GIMAP5*, which both result in T cell deficiency when knocked out in mice^34^, were significantly downregulated (adj p < 0.05, log2(fold change) < -0.5) in each comparison, except for *GIMAP5* in *TET2* CD8+T cells which had a p-value of 0.235.

**Fig. 5:**
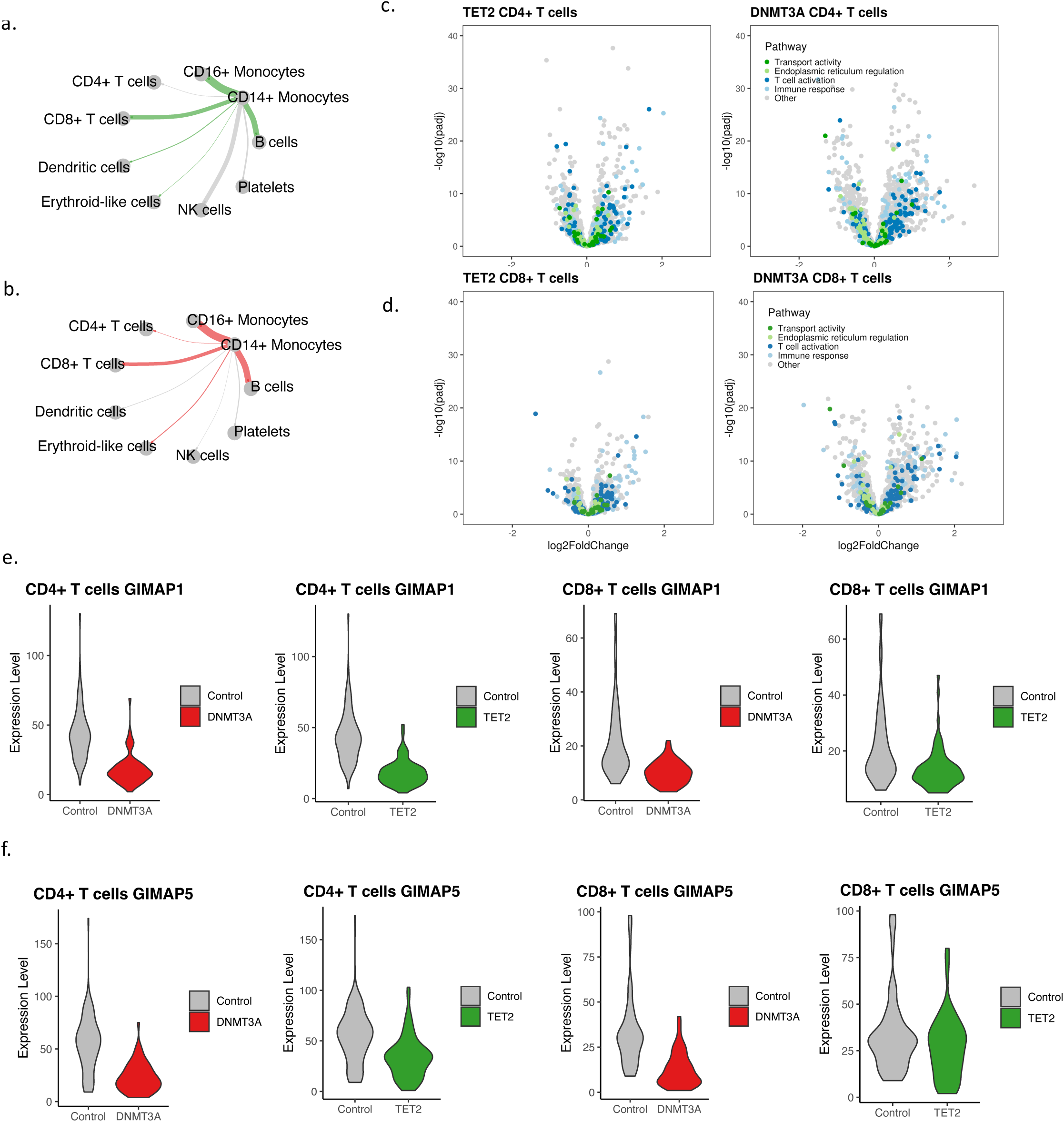
Genotype-grouped CD4+ and CD8+ T cell vs control comparisons highlight alterations to T cell activation and differentiation in CH. **A)** Circle plot displaying predicted differential interaction strength between CD14+ monocytes and relevant hematopoietic cells from *TET2* samples compared to controls, as determined by CellChat. Line thickness corresponds to differential interaction strength. Green color indicates increased signaling in *TET2* samples compared to controls. Grey indicates decreased signaling. **B)** Circle plot displaying predicted differential interaction strength between CD14+ monocytes and relevant hematopoietic cells from *DNMT3A* samples compared to controls, as determined by CellChat. Line thickness corresponds to differential interaction strength. Red color indicates increased signaling in *DNMT3A* samples compared to controls. Grey indicates decreased signaling. **C)** Volcano plot showing results of differential expression analysis comparing CD4+ T cells from patients with *TET2* (left) or *DNMT3A* (right) mutations to CD4+ T cells from controls, colored by biological pathway. **D)** Volcano plot showing results of differential expression analysis comparing CD8+ T cells from patients with *TET2* (left) or *DNMT3A* (right) mutations to CD8+ T cells from controls, colored by biological pathway. **E)** Violin plots displaying expression of *GIMAP1* in CD4+ T cells and CD8+ T cells from controls and from patients with CH mutations. **F)** Violin plots displaying expression of *GIMAP5* in CD4+ T cells and CD8+ T cells from controls from patients with CH mutations.

## Discussion

Here, we present transcriptional profiling and characterization of *DNMT3A* and *TET2* CH in human peripheral blood. By using a novel approach that integrates multimodal single-cell RNA sequencing with scDNA sequencing to link mitochondrial mutations to somatic nuclear mutations, we simultaneously resolve DNA mutational status and cell state. Our study revealed CH mutation specific aberrations in cellular state allowing several conclusions.

First, we identified CD14+ monocytes as drivers of CH-associated inflammation in the peripheral blood in both *TET2* and *DNMT3A* CH. Specifically, we found *TET2* CH mutant CD14+ monocytes harbored important differences suggesting cell intrinsic mechanisms are important to *TET2* phenotypes. In relation to non-mutant wildtype monocytes from *TET2* patients, mutant CD14+ monocytes exhibited significant differences across important inflammatory genes including *IL1B*, *CXCL3*, and *CXCL1*, a phenomenon that was not seen in *DNMT3A*. Furthermore, intracellular monocyte signaling via STAT3 in response to IL-6 exhibited a VAF dependent increase further supporting the notion that mutant *TET2* monocytes exhibit cell intrinsic signaling patterns. These experiments suggest a precision medicine therapeutic for *TET2* CH. A recent analysis of the Canakinumab Anti-inflammatory Thrombosis Outcomes Trial (CANTOS) found that IL-1B antagonist, Canakinumab^36^, reduced cardiovascular risk in *TET2* but not *DNMT3A* CH patients ^2,37^. Our data provides a mechanistic rationale for a genotype-specific approach to treat CH, a finding only possible with the ability to partition mutant and wildtype cells from the same sample.

Second, collective differences between CD14+ monocytes from both *TET2* and DNMT3A and controls identify novel gene targets and signaling pathways. Computationally-inferred outgoing signaling in monocytes from *TET2* and *DNMT3A* patients indicated a notable increase in *MIF* signaling. *MIF* resides as a pre-formed peptide in a variety of cell types and binds with its receptors CXCR2 as well as CXCR4 to promote the recruitment of monocytes and T cells to sites of tissue injury^38^. Recruitment of hyperinflammatory monocytes has been identified as the initiating event in the development of atherosclerotic plaques^39^. *ADGRE5* which encodes CD97 had significantly higher expression among monocytes from *DNMT3A* patients. The protein product of this gene promotes the adhesion and migration to sites of inflammation^40^ and has been associated with rheumatoid arthritis^41^. Therefore, MIF and *ADGRE5* may represent novel targets in treating inflammation associated with *TET2* and *DNMT3A* CH.

Third, our study clarified the cell-extrinsic effects of CH-mutations in peripheral blood. Comparison between T cells from CH samples and controls highlighted significant effects of CH on both T cell differentiation and T cell activation. We observed consistent downregulation of the GIMAP protein family in CD4+ and CD8+ T cells in both *TET2* and *DNMT3A* samples. Work in mice has established that knockout of GIMAP proteins impairs development of T and B cells, resulting in a relative T/B deficiency and a myeloid skew^34^, similar to what is observed in CH. GIMAP proteins are regulated together under the direction of the transcription factors RUNX1, GATA3, and TAL1^42,43^. DNMT3A directly binds to RUNX1 and GATA3^42^, and TAL1 expression has been shown to be disrupted by knockout of both *TET2* and *DNMT3A*^44^. Further work investigating the effect of *TET2* and *DNMT3A* mutations on GIMAP expression and subsequent differentiation is warranted.

Our study has several limitations. First, while *TET2* and *DNMT3A* mutations make up approximately 2/3 of all CH mutations, CH represents a diverse set of mutations in >70 genes. These CH mutations are likely to have divergent effects from those we describe here. We also binned samples with co-occurring mutations additional to *TET2* or *DNMT3A* mutations to increase our sample set size, though this may introduce a source of variability. Second, we cannot exclude that CH with small clones below our limit of detection are present in our control samples. However, we would expect minimal pathological effect given the marginal size of these clones. Third, a shortcoming of our work is the absence of neutrophils in PBMC samples. Given the myeloid bias of CH mutations, it is likely that neutrophils also harbor mutations, and so their functional consequences within the periphery require investigation. Finally, two of our *TET2* samples have co-occurring *SRSF2* mutations, which, as others have noted, are unfortunately not able to be detected consistently with scDNA-seq. Based on VAF data, these mutations are subclonal to TET2 in each case, this is due to GC content in the region^45^; thus, we are unable to assess the effect of this mutation our samples on a single cell level. Future efforts to improve single cell genotyping of this gene may help elucidate this. Overall, our study provides mechanistic support for a genotype specific precision medicine approach for future CH therapeutics.

## Data and materials availability

All filtered count matrices and DGE tables are available on Open Science Framework at https://osf.io/rac5w/. Seurat objects will be made available through the Chan Zuckerberg Initiative database. All data analysis was completed using R (v4.1.2) on the Terra.bio cloud platform. All R files used to generate the figures and tables will be made publicly available on GitHub upon publication and can be made available for reviewers upon request.

## Acknowledgements

We thank Angela Jones for her assistance with sequencing efforts associated with this work. A.G.B. is supported by a Burroughs Wellcome Foundation Career Award for Medical Scientists and the NIH Director’s Early Independence Award (DP5-OD029586). P.B.F is supported by a NIH K23HL138291 and a Mark Foundation Endeavor Award. P.v.G. is supported by the Ludwig Center at Harvard, the NIH (R00CA218832), Gilead Sciences, the Bertarelli Rare Cancers Fund, the Starr Cancer Consortium, the William Guy Forbeck Research Foundation, and is an awardee of the Glenn Foundation for Medical Research and American Federation for Aging Research (AFAR) Grant for Junior Faculty. M.R.S. is supported by NIH 1R01CA262287 and 1U01OH012271, an LLS Clinical Scholar Award, the Biff Ruttenburg Foundation, the Adventure Alle Fund, the Beverly and George Rawlings Research Directorship, the E.P. Evans MDS Foundation.

## Author contributions

J.B.H. and P.B. designed the study, facilitated data collection, conducted formal analysis and interpretation of results, generated figures, prepared the original draft and edited the manuscript.

A.C.P., C.V., and M.T.J. collected data, conducted formal analysis and interpretation of results, generated figures, prepared the original draft and edited the manuscript.

J.U., C.R.P., S.O., and N.N.H. facilitated sample curation and data collection.

B.S. and A.A. provided analysis software.

P.V.G. provided resources, analysis software, and edited manuscript.

A.J.S. and M.R.S. facilitated sample curation, provided resources and project administration, and edited the manuscript.

A.G.B. and P.B.F. conceived and supervised the study, provided funding for the study, provided resources and project administration, conducted formal analysis and interpretation of results, generated figures, prepared the original draft and edited the manuscript.

## Competing Interests Statement

All unrelated to the present work: M.R.S. reports personal fees from AbbVie, BMS, CTI, Sierra Oncology, Novartis, grants from Astex and Incyte, personal fees and other support from Karyopharm, Ryvu, personal fees from Sierra Oncology, grants and personal fees from Takeda, and TG Therapeutics outside the submitted work. P.B.F. reports research funding from Novartis. A.G.B. is a scientific co-founder and has equity in TenSixteen Bio. All other authors declare that they have no competing interests.

Supplementary Information is available for this paper.

Correspondence and requests for materials should be addressed to: Alexander G. Bick, P. Brent Ferrell

