## Supplement for "Multiomic Profiling of Human Clonal Hematopoiesis Reveals Genotype and Cell-Specific Inflammatory Pathway Activation"

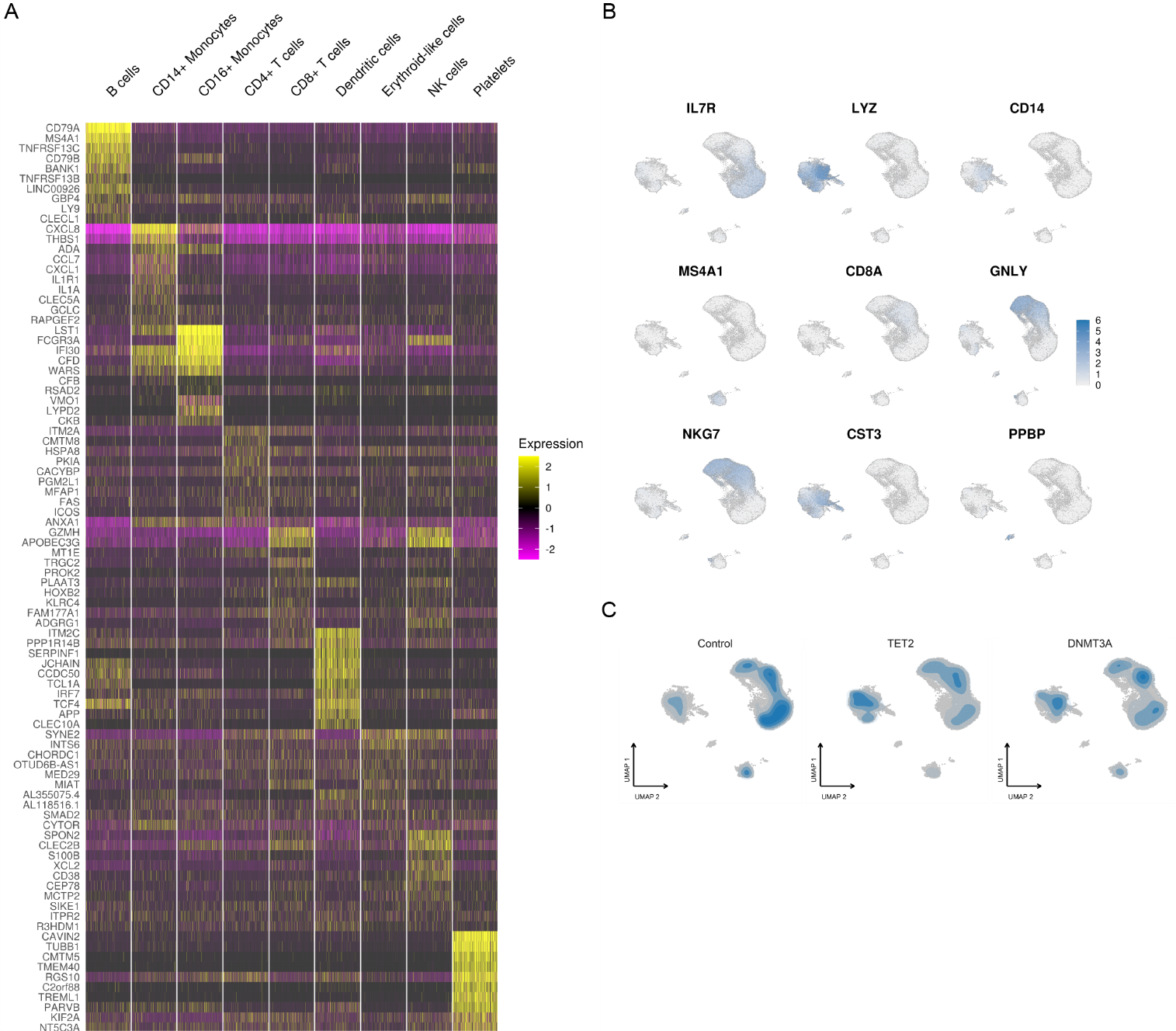

**Supplemental Figure 1: Cell type assignments align with expectations from canonical markers.** A) Heatmap displaying expression of canonical markers for each cell type. B) Feature plots showing expression of canonical markers in UMAP space. C) UMAPs depicting cell type clusters for control (n = 34,774), *TET2* (n = 41,260), and *DNMT3A* patients (n = 28,944) with density of points indicated in blue.

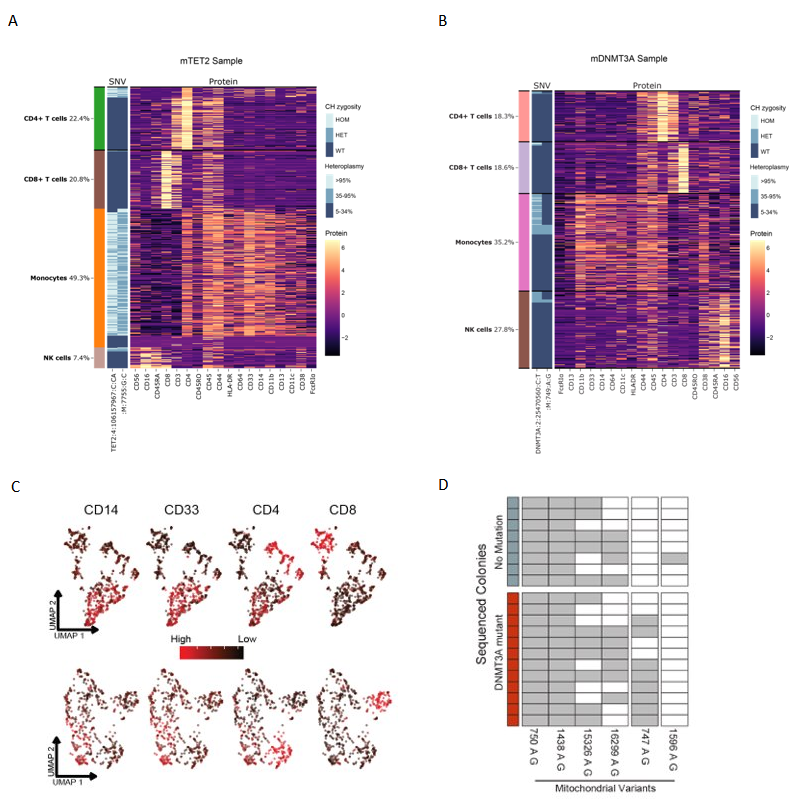

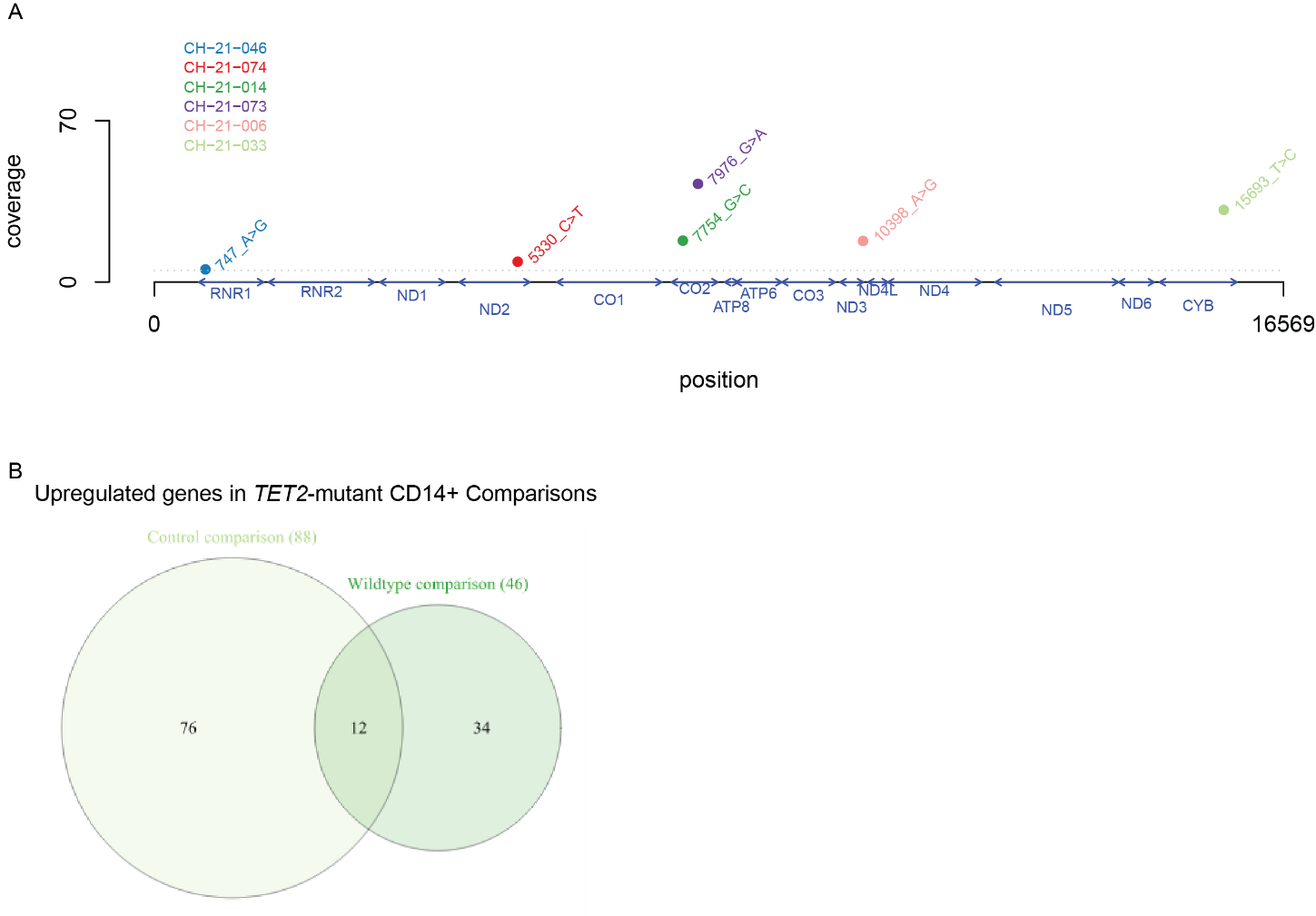

**Supplemental Figure 2: Single-cell DNA-sequencing reveals concordance of mitochondrial variants and CHIP mutations in A) *TET2*-mutant sample (n=999 cells) and B) *DNMT3A*-mutant sample (n=665 cells)**. Heatmap highlighting hierarchical clustering of AOCs showing CLR-normalized counts. Columns represent AOCs and rows represent cells. The cell barcodes are sorted by genotype status (presence or absence of variant, based on zygosity). C) Expression of select AOCs in UMAP space in *TET2*-mutant sample (above) and *DNMT3A*-mutant sample (below). D) Selection strategy for mitochondrial variants. Matrix highlighting the presence of specific mitochondrial mutations identified in individual colonies from the methylcellulose assay. Rows represent individual colonies and CH mutation status is noted by color. Columns represent mitochondrial mutations. Shading notes presence of a mitochondrial mutation in a specific colony. AOC - antibody oligonucleotide conjugates PBMC – peripheral blood mononuclear cells CB – Cell Barcode HET – heterozygous call by clustering algorithm HOM – homozygous call by clustering algorithm WT – wildtype call by clustering algorithm.

**Supplemental Figure 3: Mitochondrial variant calling and mutant cell differential gene expression comparisons.** A) Coverage plot of selected mitochondrial variants from individual CH samples (*TET2*-mutant, n=4; *DNMT3A*-mutant, n=2). Variants with coverage greater than 5 were selected for downstream analysis. B) Overlap of significantly upregulated genes in *TET2*-mutant cells compared to wildtype or control sample cells. 88 genes and 46 genes were upregulated in the control and wildtype comparisons, respectively. 12 genes were shared between the comparisons.

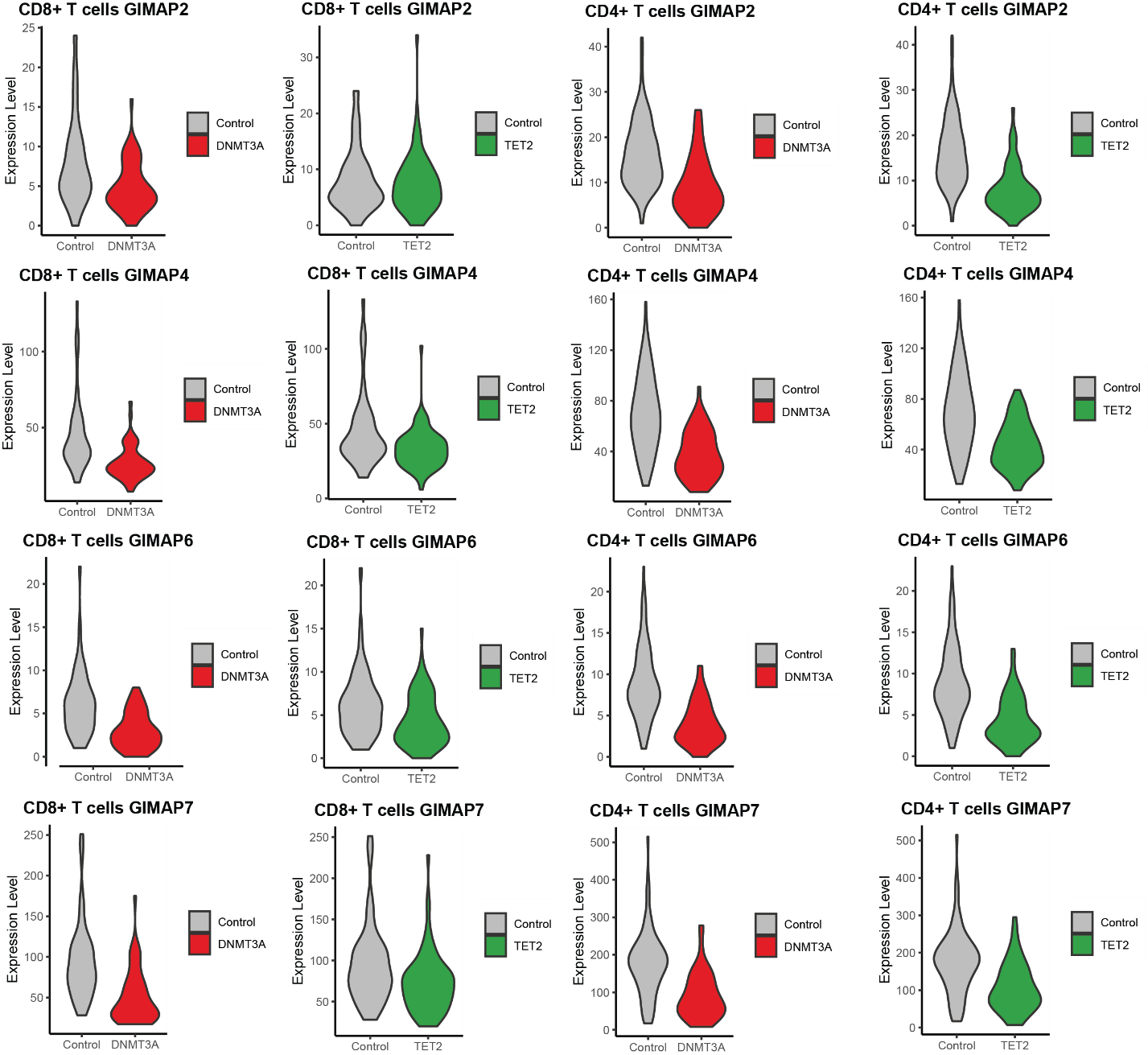

**Supplemental Figure 4: Differential expression analysis reveals consistent downregulation of GIMAP genes in T cells from CH patients.** Violin plots showing expression of GIMAP genes in CD4+ and CD8+ T cells from CH patients and from controls, based on metacell aggregation.

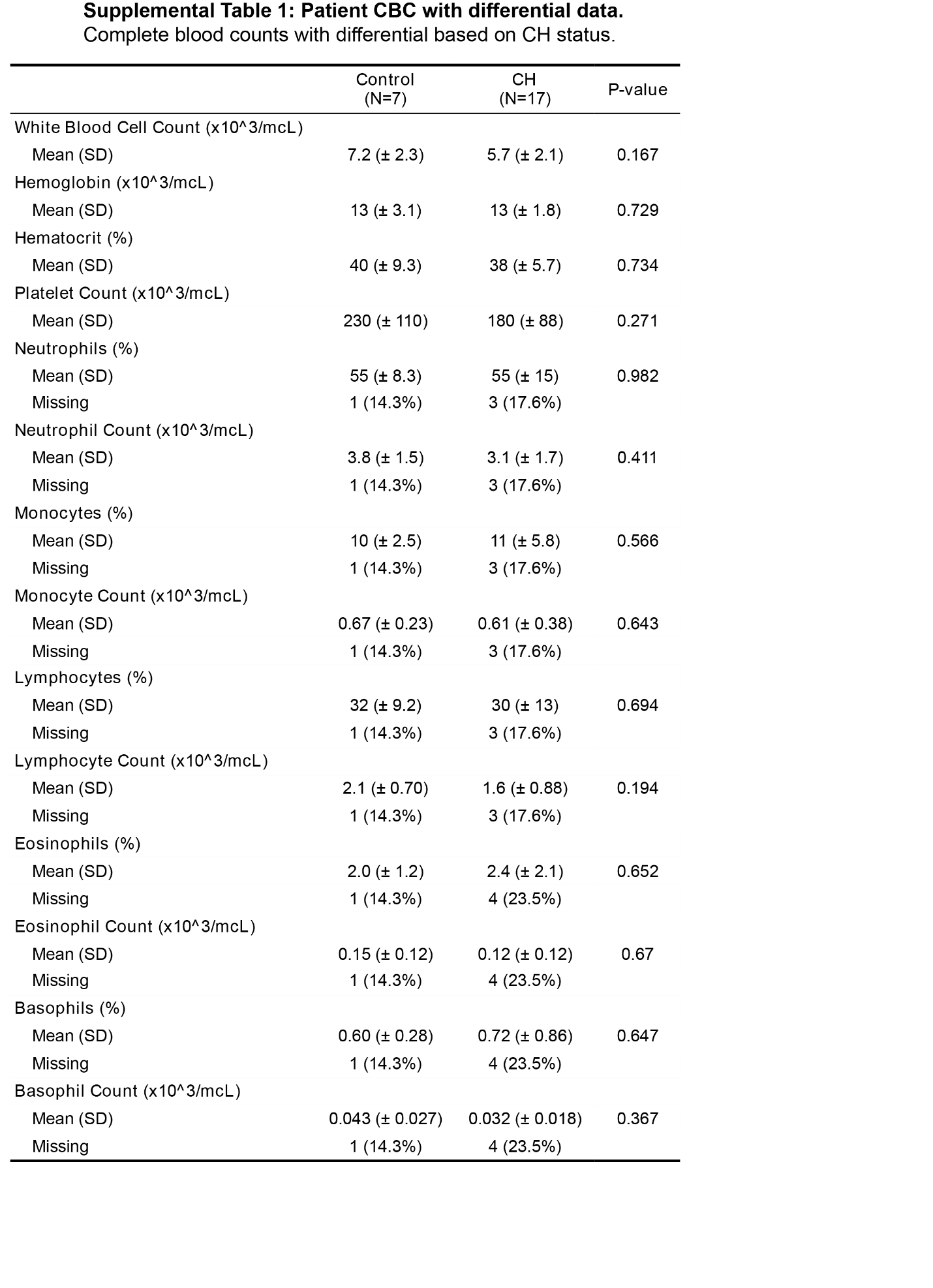

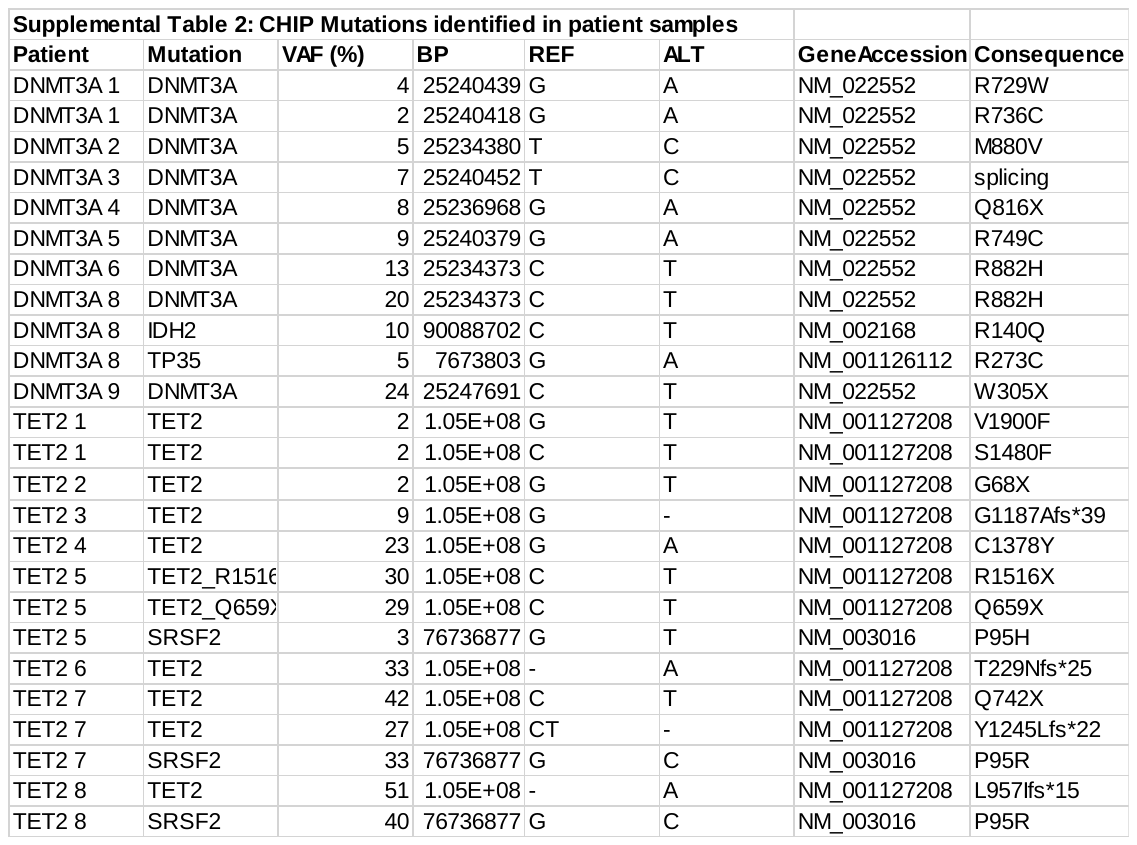

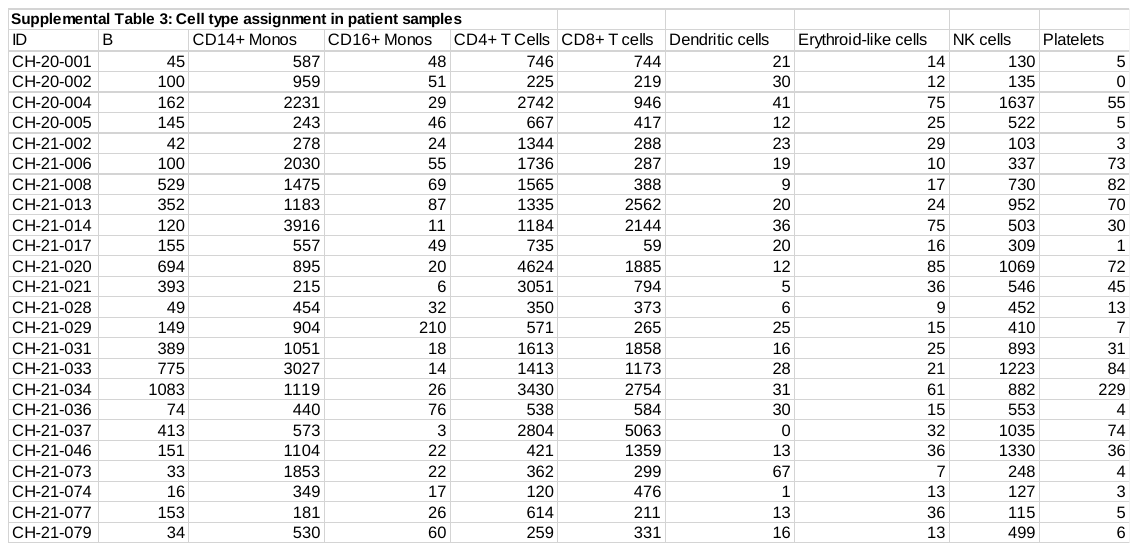

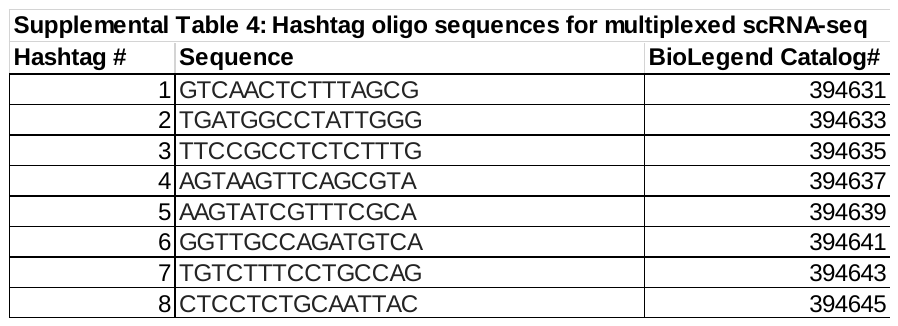

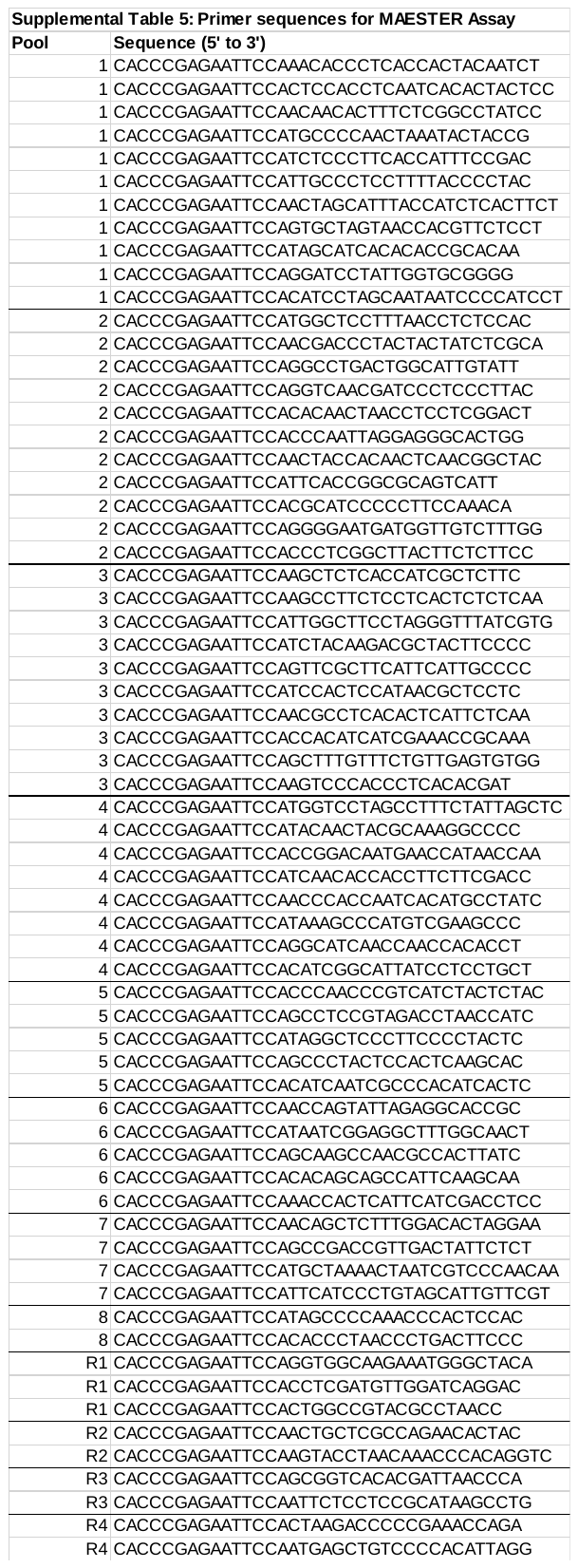

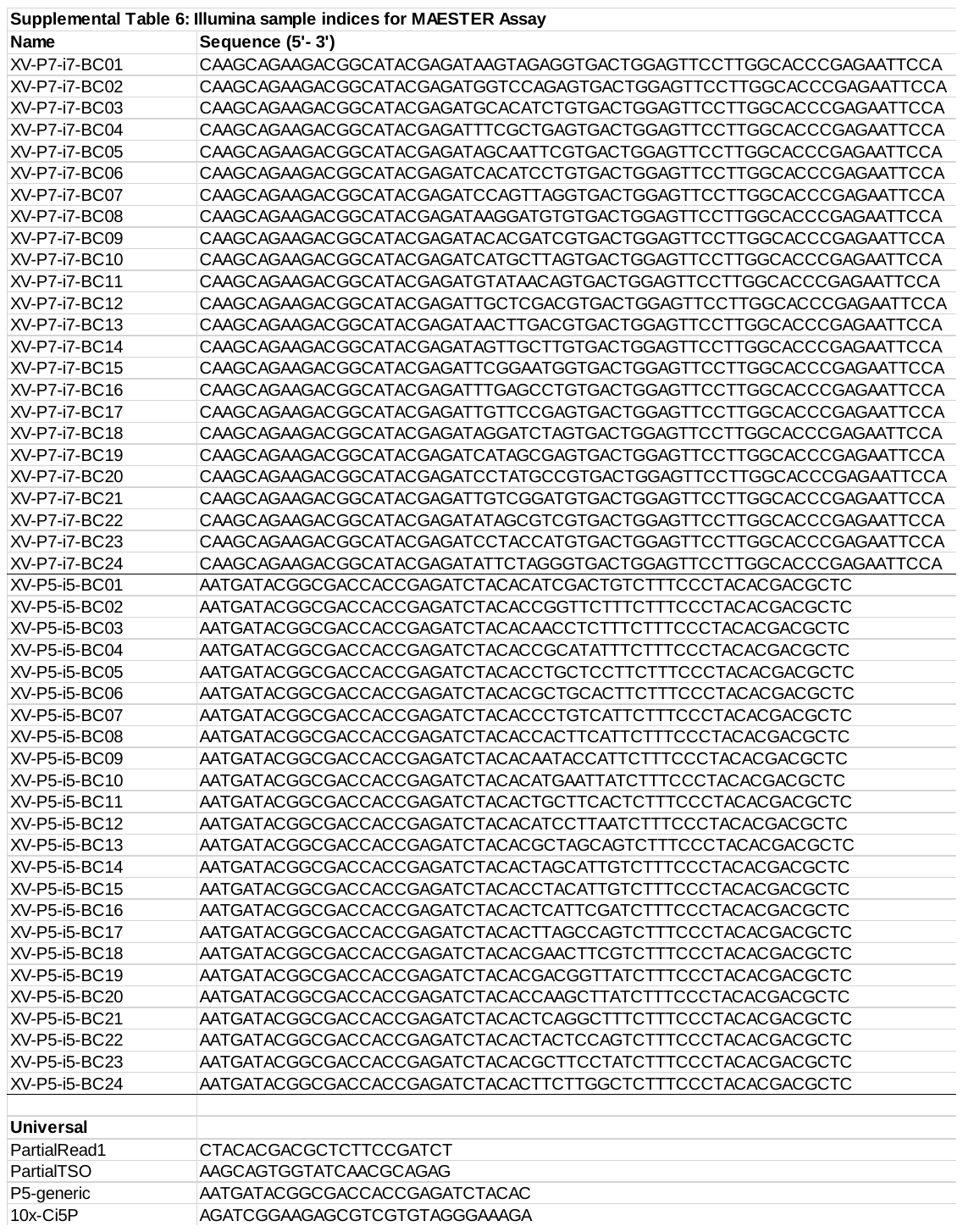

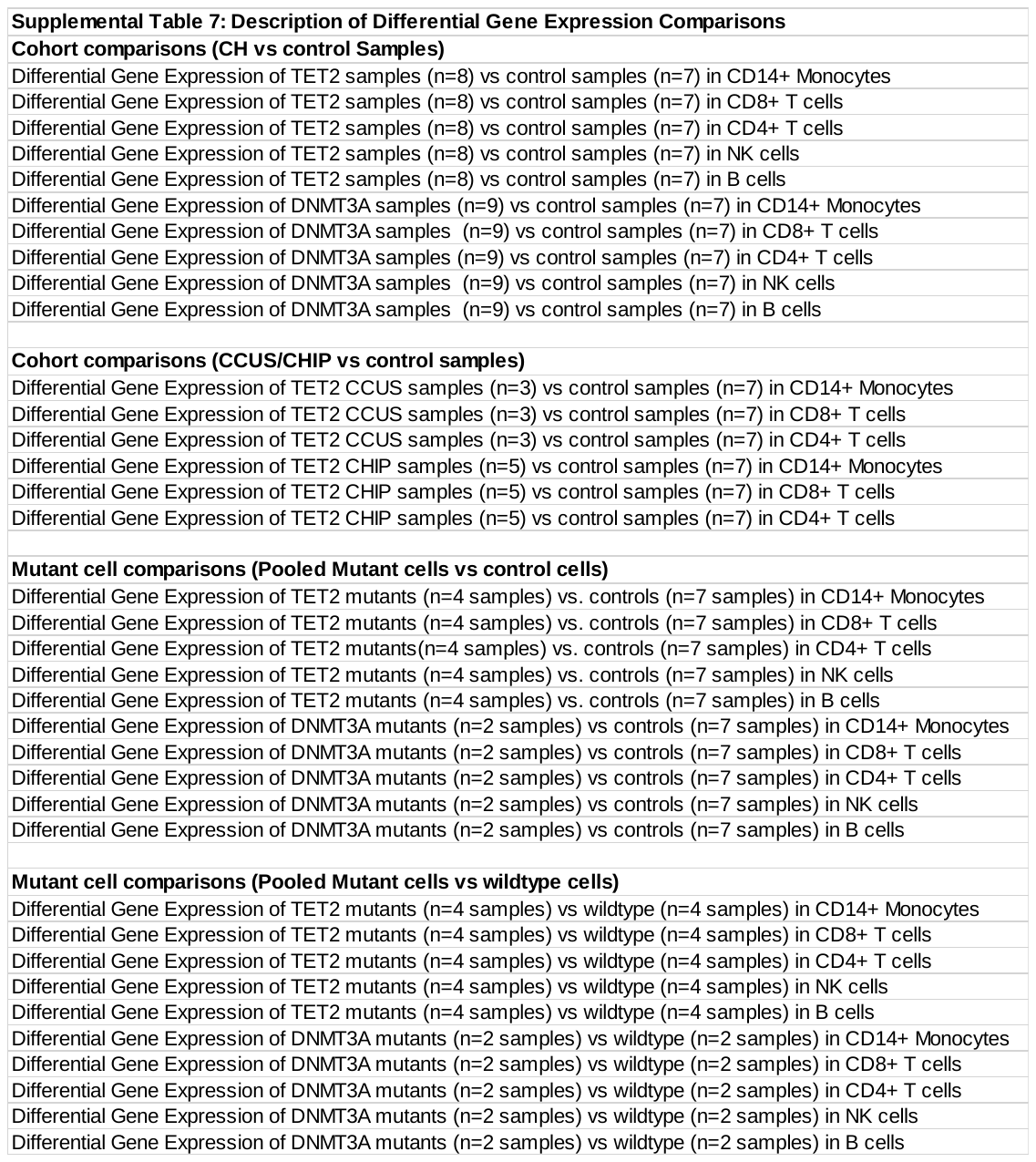

**Supplemental Table 8: Amplicon Locations for Tapestri protocol**

Chromosome	Start	End	ID

chr1	115256488	115256723	AML_v2_NRAS_115256512

chr1	115258610	115258825	AML_v2_NRAS_115258635

chr2	25457144	25457372	AML_v2_DNMT3A_25457166

chr2	25458519	25458763	AML_v2_DNMT3A_25458540

chr2	25459794	25460046	AML_v2_DNMT3A_25459813

chr2	25461881	25462137	AML_v2_DNMT3A_25461902

chr2	25463107	25463346	AML_v2_DNMT3A_25463127

chr2	25463494	25463717	AML_v2_DNMT3A_25463515

chr2	25464422	25464618	AML_v2_DNMT3A_25464443

chr2	25466622	25466871	AML_v2_DNMT3A_25466642

chr2	25467013	25467220	AML_v2_DNMT3A_25467033

chr2	25467372	25467631	AML_v2_DNMT3A_25467391

chr2	25468111	25468332	AML_v2_DNMT3A_25468130

chr2	25469007	25469231	AML_v2_DNMT3A_25469026

chr2	25469408	25469606	AML_v2_DNMT3A_25469428

chr2	25469926	25470185	AML_v2_DNMT3A_25469945

chr2	25470404	25470663	AML_v2_DNMT3A_25470423

chr2	25470931	25471190	AML_v2_DNMT3A_25470951

chr2	25472516	25472728	AML_v2_DNMT3A_25472538

chr2	198266104	198266322	AML_v2_SF3B1_198266130

chr2	198266443	198266688	AML_v2_SF3B1_198266472

chr2	198266694	198266913	AML_v2_SF3B1_198266718

chr2	198267320	198267569	AML_v2_SF3B1_198267340

chr2	209113086	209113297	AML_v2_IDH1_209113110

chr3	128200078	128200327	AML_v2_GATA2_128200102

chr3	128200669	128200928	AML_v2_GATA2_128200689

chr3	128202700	128202899	AML_v2_GATA2_128202719

chr4	55561580	55561792	AML_v2_KIT_55561605

chr4	55569876	55570095	AML_v2_KIT_55569899

chr4	55592078	55592282	AML_v2_KIT_55592099

chr4	55593510	55593744	AML_v2_KIT_55593532

chr4	55593964	55594183	AML_v2_KIT_55593987

chr4	55599271	55599486	AML_v2_KIT_55599293

chr4	55602652	55602862	AML_v2_KIT_55602675

chr4	106154924	106155158	AML_v2_TET2_106154951

chr4	106155159	106155416	AML_v2_TET2_106155178

chr4	106155470	106155729	AML_v2_TET2_106155489

chr4	106155914	106156173	AML_v2_TET2_106155932

chr4	106156238	106156489	AML_v2_TET2_106156259

chr4	106156504	106156762	AML_v2_TET2_106156523

chr4	106156794	106157046	AML_v2_TET2_106156812

chr4	106157078	106157332	AML_v2_TET2_106157098

chr4	106157427	106157679	AML_v2_TET2_106157447

chr4	106157757	106158015	AML_v2_TET2_106157777

chr4	106158030	106158288	AML_v2_TET2_106158049

chr4	106158294	106158544	AML_v2_TET2_106158314

chr4	106158546	106158805	AML_v2_TET2_106158576

chr4	106162389	106162618	AML_v2_TET2_106162415

chr4	106163953	106164192	AML_v2_TET2_106163979

chr4	106164706	106164952	AML_v2_TET2_106164725

chr4	106180719	106180956	AML_v2_TET2_106180745

chr4	106182830	106183072	AML_v2_TET2_106182858

chr4	106190734	106190955	AML_v2_TET2_106190758

chr4	106193541	106193777	AML_v2_TET2_106193572

chr4	106193778	106194032	AML_v2_TET2_106193797

chr4	106194036	106194295	AML_v2_TET2_106194057

chr4	106196180	106196429	AML_v2_TET2_106196202

chr4	106196438	106196692	AML_v2_TET2_106196456

chr4	106196772	106197024	AML_v2_TET2_106196791

chr4	106197029	106197278	AML_v2_TET2_106197048

chr4	106197336	106197593	AML_v2_TET2_106197355

chr4	106197599	106197858	AML_v2_TET2_106197618

chr5	170837385	170837659	AML_v2_NPM1_170837412

chr7	148504722	148504971	AML_v2_EZH2_148504743

chr7	148506026	148506265	AML_v2_EZH2_148506050

chr7	148506372	148506589	AML_v2_EZH2_148506394

chr7	148507405	148507618	AML_v2_EZH2_148507427

chr7	148508697	148508930	AML_v2_EZH2_148508719

chr7	148511033	148511276	AML_v2_EZH2_148511054

chr7	148511991	148512230	AML_v2_EZH2_148512017

chr7	148514918	148515124	AML_v2_EZH2_148514941

chr7	148523627	148523866	AML_v2_EZH2_148523652

chr7	148525654	148525888	AML_v2_EZH2_148525675

chr7	148526738	148526948	AML_v2_EZH2_148526763

chr7	148529631	148529890	AML_v2_EZH2_148529658

chr7	148543468	148543693	AML_v2_EZH2_148543492

chr7	148544267	148544493	AML_v2_EZH2_148544293

chr9	5073699	5073902	AML_v2_JAK2_5073725

chr11	32413428	32413633	AML_v2_WT1_32413452

chr11	32414186	32414405	AML_v2_WT1_32414209

chr11	32417759	32417989	AML_v2_WT1_32417780

chr11	32421512	32421750	AML_v2_WT1_32421532

chr11	32439083	32439321	AML_v2_WT1_32439105

chr12	25378535	25378794	AML_v2_KRAS_25378559

chr12	25380239	25380478	AML_v2_KRAS_25380260

chr12	25398207	25398433	AML_v2_KRAS_25398232

chr12	112888116	112888350	AML_v2_PTPN11_112888140

chr12	112890995	112891234	AML_v2_PTPN11_112891019

chr12	112910668	112910907	AML_v2_PTPN11_112910689

chr12	112915378	112915582	AML_v2_PTPN11_112915401

chr12	112924189	112924405	AML_v2_PTPN11_112924215

chr12	112926204	112926424	AML_v2_PTPN11_112926226

chr12	112926825	112927050	AML_v2_PTPN11_112926847

chr13	28589757	28589977	AML_v2_FLT3_28589783

chr13	28592474	28592726	AML_v2_FLT3_28592494

chr13	28597498	28597727	AML_v2_FLT3_28597520

chr13	28601131	28601358	AML_v2_FLT3_28601153

chr13	28602303	28602559	AML_v2_FLT3_28602324

chr13	28608189	28608395	AML_v2_FLT3_28608210

chr13	28608474	28608696	AML_v2_FLT3_28608497

chr13	28609572	28609790	AML_v2_FLT3_28609594

chr13	28610015	28610260	AML_v2_FLT3_28610043

chr15	90631741	90631990	AML_v2_IDH2_90631760

chr17	7572907	7573129	AML_v2_TP53_7572930

chr17	7573974	7574178	AML_v2_TP53_7573996

chr17	7576760	7576976	AML_v2_TP53_7576782

chr17	7577015	7577264	AML_v2_TP53_7577035

chr17	7577398	7577636	AML_v2_TP53_7577424

chr17	7578076	7578315	AML_v2_TP53_7578098

chr17	7578363	7578618	AML_v2_TP53_7578383

chr17	7579859	7580118	AML_v2_TP53_7579878

chr17	74732192	74732450	AML_v2_SRSF2_74732219

chr20	30956750	30956969	AML_v2_ASXL1_30956774

chr20	31015814	31016051	AML_v2_ASXL1_31015840

chr20	31021138	31021366	AML_v2_ASXL1_31021160

chr20	31021439	31021659	AML_v2_ASXL1_31021460

chr20	31022169	31022417	AML_v2_ASXL1_31022192

chr20	31022720	31022978	AML_v2_ASXL1_31022741

chr20	31023009	31023248	AML_v2_ASXL1_31023032

chr20	31023262	31023492	AML_v2_ASXL1_31023285

chr20	31023557	31023761	AML_v2_ASXL1_31023578

chr21	36171568	36171811	AML_v2_RUNX1_36171592

chr21	36206684	36206913	AML_v2_RUNX1_36206703

chr21	36231693	36231937	AML_v2_RUNX1_36231714

chr21	36252820	36253046	AML_v2_RUNX1_36252844

chr21	44514550	44514808	AML_v2_U2AF1_44514570

chr21	44524417	44524634	AML_v2_U2AF1_44524438

chrM	13	261	AMPL538859

chrM	262	524	AMPL538860

chrM	525	774	AMPL538861

chrM	775	1023	AMPL538862

chrM	1024	1272	AMPL538863

chrM	1273	1468	AMPL538864

chrM	1469	1695	AMPL538865

chrM	1696	1936	AMPL538866

chrM	1938	2186	AMPL538867

chrM	2187	2431	AMPL538868

chrM	2432	2630	AMPL538869

chrM	2631	2824	AMPL538870

chrM	2825	3076	AMPL538871

chrM	3078	3320	AMPL538872

chrM	3321	3548	AMPL538873

chrM	3549	3797	AMPL538874

chrM	3798	4003	AMPL538875

chrM	4011	4260	AMPL538876

chrM	4284	4548	AMPL538877

chrM	4646	4895	AMPL538878

chrM	4908	5157	AMPL538879

chrM	5190	5439	AMPL538880

chrM	5440	5689	AMPL538881

chrM	5725	5983	AMPL538882

chrM	5987	6248	AMPL538883

chrM	6249	6494	AMPL538884

chrM	6495	6741	AMPL538885

chrM	6742	6966	AMPL538886

chrM	6967	7201	AMPL538887

chrM	7202	7450	AMPL538888

chrM	7451	7684	AMPL538889

chrM	7685	7923	AMPL538890

chrM	7924	8153	AMPL538891

chrM	8154	8389	AMPL538892

chrM	8391	8645	AMPL538893

chrM	8646	8912	AMPL538894

chrM	8913	9160	AMPL538895

chrM	9163	9416	AMPL538896

chrM	9417	9669	AMPL538897

chrM	9670	9916	AMPL538898

chrM	9919	10155	AMPL538899

chrM	10156	10384	AMPL538900

chrM	10387	10637	AMPL538901

chrM	10663	10913	AMPL538902

chrM	10914	11153	AMPL538903

chrM	11155	11401	AMPL538904

chrM	11402	11640	AMPL538905

chrM	11642	11869	AMPL538906

chrM	11870	12118	AMPL538907

chrM	12119	12366	AMPL538908

chrM	12367	12556	AMPL538909

chrM	12557	12755	AMPL538910

chrM	12756	12970	AMPL538911

chrM	12971	13208	AMPL538912

chrM	13209	13458	AMPL538913

chrM	13462	13710	AMPL538914

chrM	13711	13956	AMPL538915

chrM	13957	14206	AMPL538916

chrM	14207	14401	AMPL538917

chrM	14402	14639	AMPL538918

chrM	14640	14861	AMPL538919

chrM	14862	15109	AMPL538920

chrM	15110	15351	AMPL538921

chrM	15352	15595	AMPL538922

chrM	15596	15843	AMPL538923

chrM	15844	16093	AMPL538924

chrM	16097	16346	AMPL538925

chrM	16347	16569	AMPL538926

Supplemental Table 9

Summary Statistics for Violin Plots

| **Figure** | **Celltype** | **Gene** | **Gneotype** | **Mean** | **SD** |
| --- | --- | --- | --- | --- | --- |
| 2A | CD14 Monocytes | IL1B | TET2 | 74.89 | 1.08757 |
| 2A | CD14 Monocytes | IL1B | Control | 45.01 | 1.464576 |
| 2A | CD14 Monocytes | CXCL3 | TET2 | 28.96 | 1.712802 |
| 2A | CD14 Monocytes | CXCL3 | Control | 14.86 | 1.585621 |
| 2A | CD14 Monocytes | CXCL1 | TET2 | 4.71 | 1.197598 |
| 2A | CD14 Monocytes | CXCL1 | Control | 1.97 | 0.8849939 |
| 2B | CD14 Monocytes | CCL4 | DNMT3A | 53.5 | 1.055184 |
| 2B | CD14 Monocytes | CCL4 | Control | 34.34 | 1.769285 |
| 2B | CD14 Monocytes | CCL2 | DNMT3A | 16.46 | 1.462901 |
| 2B | CD14 Monocytes | CCL2 | Control | 10.92 | 1.667394 |
| 2B | CD14 Monocytes | CCL7 | DNMT3A | 4.15 | 1.175376 |
| 2B | CD14 Monocytes | CCL7 | Control | 3.16 | 1.010029 |
| 4A | CD14+ Monocytes | FN1 | Control | 0.9464286 | 1.4575152 |
| 4A | CD14+ Monocytes | FN1 | TET2 | 12.5492958 | 16.3791848 |
| 4A | CD14+ Monocytes | FCER2 | Control | 0.5178571 | 0.8736771 |
| 4A | CD14+ Monocytes | FCER2 | TET2 | 7.6478873 | 13.3709737 |
| 4B | CD14+ Monocytes | IFITM2 | Control | 33.6428571 | 30.2200587 |
| 4B | CD14+ Monocytes | IFITM2 | DNMT3A | 77.5777778 | 53.4484899 |
| 4B | CD14+ Monocytes | ADGRE5 | Control | 26.3035714 | 17.0431747 |
| 4B | CD14+ Monocytes | ADGRE5 | DNMT3A | 40.8222222 | 21.2140391 |
| 5E | CD4+ T cells | GIMAP1 | Control | 43.6990291 | 18.1160935 |
| 5E | CD4+ T cells | GIMAP1 | TET2 | 18.5285714 | 8.9777849 |
| 5E | CD4+ T cells | GIMAP1 | DNMT3A | 18.1969697 | 10.9617793 |
| 5E | CD8+ T cells | GIMAP1 | Control | 21.875 | 14.1499277 |
| 5E | CD8+ T cells | GIMAP1 | TET2 | 13.4810127 | 7.2197393 |
| 5E | CD8+ T cells | GIMAP1 | DNMT3A | 10.1296296 | 4.2428466 |
| 5E | CD4+ T cells | GIMAP5 | Control | 54.5728155 | 26.2923242 |
| 5E | CD4+ T cells | GIMAP5 | TET2 | 33.8857143 | 20.5655846 |
| 5E | CD4+ T cells | GIMAP5 | DNMT3A | 23.030303 | 13.6200866 |
| 5E | CD8+ T cells | GIMAP5 | Control | 33.5178571 | 20.4548203 |
| 5E | CD8+ T cells | GIMAP5 | TET2 | 27.0253165 | 16.8728083 |
| 5E | CD8+ T cells | GIMAP5 | DNMT3A | 12.6481481 | 9.4970848 |
| Supp 4 | CD4+ T cells | GIMAP2 | Control | 16.2038835 | 7.206151 |
| Supp 4 | CD4+ T cells | GIMAP2 | TET2 | 8 | 5.0389785 |
| Supp 4 | CD4+ T cells | GIMAP2 | DNMT3A | 9.1212121 | 6.2867753 |
| Supp 4 | CD8+ T cells | GIMAP2 | Control | 7.9821429 | 5.4622375 |
| Supp 4 | CD8+ T cells | GIMAP2 | TET2 | 7.4683544 | 5.1486504 |
| Supp 4 | CD8+ T cells | GIMAP2 | DNMT3A | 4.9444444 | 3.0801658 |
| Supp 4 | CD4+ T cells | GIMAP4 | Control | 70 | 29.7802409 |
| Supp 4 | CD4+ T cells | GIMAP4 | TET2 | 43.2 | 17.9319163 |
| Supp 4 | CD4+ T cells | GIMAP4 | DNMT3A | 37.6363636 | 18.3165636 |
| Supp 4 | CD8+ T cells | GIMAP4 | Control | 45.1785714 | 24.7379775 |
| Supp 4 | CD8+ T cells | GIMAP4 | TET2 | 33.4683544 | 13.1917179 |
| Supp 4 | CD8+ T cells | GIMAP4 | DNMT3A | 27.0740741 | 11.2398413 |
| Supp 4 | CD4+ T cells | GIMAP6 | Control | 8.9126214 | 4.2451975 |
| Supp 4 | CD4+ T cells | GIMAP6 | TET2 | 4.2714286 | 2.7867254 |
| Supp 4 | CD4+ T cells | GIMAP6 | DNMT3A | 3.969697 | 2.5778327 |
| Supp 4 | CD8+ T cells | GIMAP6 | Control | 6.1607143 | 3.7357087 |
| Supp 4 | CD8+ T cells | GIMAP6 | TET2 | 4.3544304 | 2.9353634 |
| Supp 4 | CD8+ T cells | GIMAP6 | DNMT3A | 3 | 2.0649821 |
| Supp 4 | CD4+ T cells | GIMAP7 | Control | 175.708738 | 84.5657455 |
| Supp 4 | CD4+ T cells | GIMAP7 | TET2 | 106.985714 | 62.9079136 |
| Supp 4 | CD4+ T cells | GIMAP7 | DNMT3A | 90.9090909 | 60.8685159 |
| Supp 4 | CD8+ T cells | GIMAP7 | Control | 94.875 | 47.7668818 |
| Supp 4 | CD8+ T cells | GIMAP7 | TET2 | 72.3037975 | 35.7840225 |
| Supp 4 | CD8+ T cells | GIMAP7 | DNMT3A | 50.5 | 32.3096749 |
